# A shark variable new antigen receptor recognizes an occluded epitope of fibroblast activation protein

**DOI:** 10.64898/2026.08.11.744296

**Authors:** Roma L. Broadberry, Kendahl L. Ott, Eric W. Lake, Tapanmitra Ravi, Colin A Hemme, Shang Chi, Jayden L. West, Gihan S. Gunaratne, Irene M. Ong, Aaron M. LeBeau, Timothy Grant

**Affiliations:** John and Jean Rowe Institute of Virology, Morgridge Institute for Research, Madison, WI 53715, USA; Department of Biochemistry, University of Wisconsin–Madison, Madison, WI 53715, USA; Department of Pathology and Laboratory Medicine, University of Wisconsin School of Medicine and Public Health, Madison, WI 53705, USA; Molecular and Cellular Pharmacology Program, University of Wisconsin School of Medicine and Public Health, Madison, WI, 53705, USA; Department of Radiology, University of Wisconsin School of Medicine and Public Health, Madison, WI 53705, USA; Department of Biostatistics and Medical Informatics, University of Wisconsin School of Medicine and Public Health, University of Wisconsin-Madison, Madison, WI, USA; University of Wisconsin Carbone Cancer Center, University of Wisconsin School of Medicine and Public Health, Madison, WI 53705, USA

**Author notes:** Corresponding Authors: **Aaron M. LeBeau**, University of Wisconsin, Madison, Wisconsin Institutes for Medical Research, 1111 Highland Avenue, Madison, WI 53705, USA., **Timothy Grant,** University of Wisconsin, Madison, Morgridge Institute for Research, 330 N. Orchard St. Madison, WI 53715, USA.

## Abstract

Variable new antigen receptors (VNARs) are the smallest naturally occurring antibody binding domains. Their size allows VNARs to access sterically restricted epitopes that are inaccessible to conventional antibodies. We recently identified a suite of VNARs that target fibroblast activation protein (FAP), a stromal serine protease indicative of extracellular matrix remodeling. The presence of FAP on the surface of cancer-associated fibroblasts (CAFs) that promote immunosuppression has made FAP a compelling therapeutic target for cancer therapy. Although antibodies targeting FAP have been developed, there is a paucity of information on how biologics engage FAP. Here, we used single-particle cryogenic electron microscopy (cryo-EM) to compare FAP recognition of three antibody architectures: a shark-derived VNAR, variable heavy (V_H_) and light domains (V_L_) of a humanized Immunoglobulin G (IgG), and a camelid-derived VHH. The humanized V_H_-V_L_ domains and camelid VHH both target a solvent-exposed β-propeller domain, whereas the VNAR binds a highly conserved, topologically recessed epitope at the FAP dimer interface. Radical-footprinting mass spectrometry (MS) further mapped two additional immune-derived VNARs to distinct FAP surfaces outside the shared β-propeller epitope. These findings demonstrate how unique VNAR architecture can expand access to underexplored FAP surfaces and establish a structural framework for rational multiepitope targeting strategies.

## Introduction

The remodeling of the extracellular matrix (ECM) is a hallmark of cancer.^1^ During tumor progression, signaling cascades activate the tumor stroma and transform fibroblasts into a reactive phenotype that can produce collagen and other ECM components.^2^ These reactive fibroblasts alter the ECM through the expression of proteases, including fibroblast activation protein (FAP). The proteolytic activity of FAP breaks down ECM proteins to drive tissue remodeling and also to activate growth factors and cytokines to further drive fibroblast activation.^3,4^ FAP is most notably expressed on the surface of cancer-associated fibroblasts (CAFs) in the stromal compartment of carcinomas.^5^ CAFs that express FAP are immunosuppressive, promote resistance to therapy, and help promote tumor growth and metastasis. Consistent with this role, FAP is expressed in the stroma of nearly 90% of all cancers of epithelial origin. In addition to cancer, FAP has been implicated in non-oncological pathologies that include fibrosis, arthritis, cardiovascular disease, and autoimmune disease.^6^

FAP is a type II transmembrane serine protease consisting of an N-terminal cytosolic domain, a transmembrane domain, and a large extracellular domain.^7^ Expressed as a 97 kDa monomer, FAP requires dimerization and glycosylation to be enzymatically active on the cell surface. As a member of the S9B prolyl oligopeptidase family, FAP is closely related to dipeptidyl peptidase 4 (DPP4), a multi-functional protease involved in glucose metabolism and immune regulation.^7,8^ FAP and DPP4 share over 50% sequence identity and adopt conserved domains: the N-terminal β-propeller domain and C-terminal α/β-hydrolase catalytic domain.^7–9^ Despite this shared fold, FAP exhibits distinct surface loops and glycosylation patterns that contribute to differences in substrate specificity and antibody recognition.^6,7^ FAP also displays endopeptidase and exopeptidase activity, distinguishing it from several related family members including DPP4 as well as DPP6, DPP8, and DPP9.^5^ Moreover, FAP is highly disease specific with little to no FAP expressed in most normal adult tissues outside of contexts such as wound healing and embryogenesis. DPP4, by contrast, is ubiquitously expressed in healthy tissue.^5,6^ The restricted expression of FAP to pathological conditions suggests that its presence could be exploited for the development of targeted diagnostic agents and therapeutics.

The development of antibodies targeting FAP remains a challenge due to the high homology among the S9B prolyl oligopeptidase family members. Furthermore, structural insights into how antibodies achieve FAP-specific recognition are limited. Since antibody structure can constrain epitope accessibility, scaffold architecture may influence which FAP surfaces can be engaged. Conventional immunoglobulins (IgGs) use paired variable heavy (V_H_) and variable light (V_L_) chains that form a 50 kDa fragment of antigen binding (Fab) domain.^10^ Each Fab variable domain contains three complementarity determining regions (CDRs) which together create a broad antigen-recognition surface at the V_H_-V_L_ interface. This planar geometry can limit access to recessed, occluded, or conformationally complex epitopes.^11^ Camelid variable-heavy-heavy (VHH) single-domain antibodies provide a smaller antibody format (∼15 kDa), comprising a single variable heavy-chain domain with three CDRs, including an elongated CDR3 loop that can improve access to cryptic or concave epitopes relative to conventional antibodies.^12,13^ Shark-derived variable new antigen receptors (VNARs) offer an even smaller single-domain format.^14^ VNARs are ∼11kDa single-chain antigen recognition domains of shark antibodies, making them the smallest antibody binding domains found in nature.^14,15^ Distinct from IgGs and VHHs, VNARs possess four variable loops: CDR1, CDR3, and the hypervariable framework regions HV2 and HV4.^16^ Among these, CDR3 is the largest and exhibits substantial sequence diversity and structural flexibility.^15^ The compact architecture of VNARs, along with their protruding loop topology and structurally diverse CDR3, supports engagement of sterically restricted epitopes that are inaccessible to larger antibody recognition domains.^16–18^ Whether these scaffold-specific geometries alter epitope accessibility and specificity on FAP remains unclear.

To address this question, we used single-particle cryogenic electron microscopy (cryo-EM) to compare FAP recognition by three antibodies representing distinct scaffold architectures: the nurse shark-derived VNAR H4-Fc, the humanized IgG Fab huB12, and the camelid VHH F7-Fc. H4-Fc was prioritized because of its high affinity, cross-reactivity with murine FAP, and nonoverlapping binding profile, whereas huB12 and F7-Fc provided structurally distinct comparators with high affinity for FAP.^19–21^ Our atomic models revealed the humanized and camelid-derived antibodies converge along the solvent exposed β-propeller domain of the FAP dimer. Interestingly, the reconstruction of FAP bound to H4-Fc found that the VNAR recognized a previously unreported epitope at the highly conserved FAP dimer interface. Complementary radical-footprinting mass spectrometry (MS) of shark-derived H15-Fc and NGS2405-Fc identified additional VNAR binding regions outside the β-propeller epitope shared by huB12 and F7-Fc. Ultimately, this study advances our understanding of anti-FAP interactions and highlights distinctive features of shark VNARs that may provide a structural basis for expanding epitope access in FAP-targeted antibody design.

## Results

### Shark VNAR H4 targets a distinct epitope on FAP

Previously, we identified a panel of anti-FAP VNARs through direct immunization of a juvenile male nurse shark with FAP that were engineered onto a human Fc scaffold (H4-Fc, H15-Fc, H17-Fc, and NGS2405-Fc).^19^ In contrast to the VNARs, the humanized huB12 and the camelid-derived F7-Fc were isolated from naïve phage display libraries. HuB12 was originally identified from a naïve murine single-chain variable fragment (scFv) library and subsequently humanized, whereas F7-Fc was obtained from a high-diversity naïve camelid library.^5,20^ Unlike H4-Fc and huB12, F7-Fc did not cross-react with mFAP and was not internalized by cells expressing FAP.^20^ Prior characterization demonstrated that H4-Fc, huB12 IgG, and F7-Fc bind recombinant and cell-surface FAP *in vitro* and localize to FAP-positive xenografts *in vivo*.^5,19–22^

To map binding profiles, we performed binning experiments using biolayer interferometry (BLI). These studies showed that H4-Fc and NGS2405-Fc recognize distinct epitopes, whereas H15-Fc and H17-Fc compete for overlapping binding sites.^19^ When H17-Fc was used as the primary antibody, no binding was observed in the presence of competing H15-Fc; however, partial binding of H17-Fc was retained when H15-Fc was used as the primary antibody.^19^ Additional binning experiments revealed that huB12 and F7-Fc share a common epitope on FAP but do not interfere with binding of the shark-derived VNAR-Fc constructs (**Supplementary Fig. 1A-B**). The humanized anti-FAP antibody sibrotuzumab, derived from the murine antibody F19, selectively targets FAP but exhibits limited internalization and showed no measurable therapeutic activity in an early phase II trial of metastatic colorectal cancer.^23,24^ In our assays, sibrotuzumab blocked binding of H17-Fc and reduced binding of H15-Fc, consistent with a shared epitope, while having no effect on the binding of huB12, F7-Fc, and NGS2405-Fc (**Supplementary Fig. 1C**).

Next, we determined single-particle cryo-EM reconstructions of purified FAP in complex with shark-derived H4-Fc, humanized huB12, and camelid-derived F7-Fc to map their respective epitopes and to compare specific mechanisms of binding. Final maps were obtained at global resolutions of 3.0 Å (EMD-78501/PDB: 37US), 2.4 Å (EMD-78502/PDB: 37UT), and 2.5 Å (EMD-78503/PDB: 37UU), respectively (**Supplemental Table 1**). In all structures, antibody fragments bound each protomer of the FAP dimer in a 1:1 stoichiometry (**Fig. 1**), while the Fc regions were not resolved due to conformational flexibility. Surface mapping of the interfaces revealed that H4 engages a recessed groove at the FAP dimer interface (**Fig. 1a-b**), in contrast to huB12 and F7 which both bind a solvent-exposed peripheral blade of the β-propeller domain (**Fig. 1a, c-d**). Consistent with these binding modes, the buried surface area (BSA) measuring the area of FAP surface buried upon complex formation, was largest for H4 (∼927 Å² per protomer) (**Fig. 1a**), followed closely by huB12 (∼884 Å²) (**Fig.1b**), and smallest for F7 (∼530 Å²) (**Fig. 1c**).

**Figure 1.**
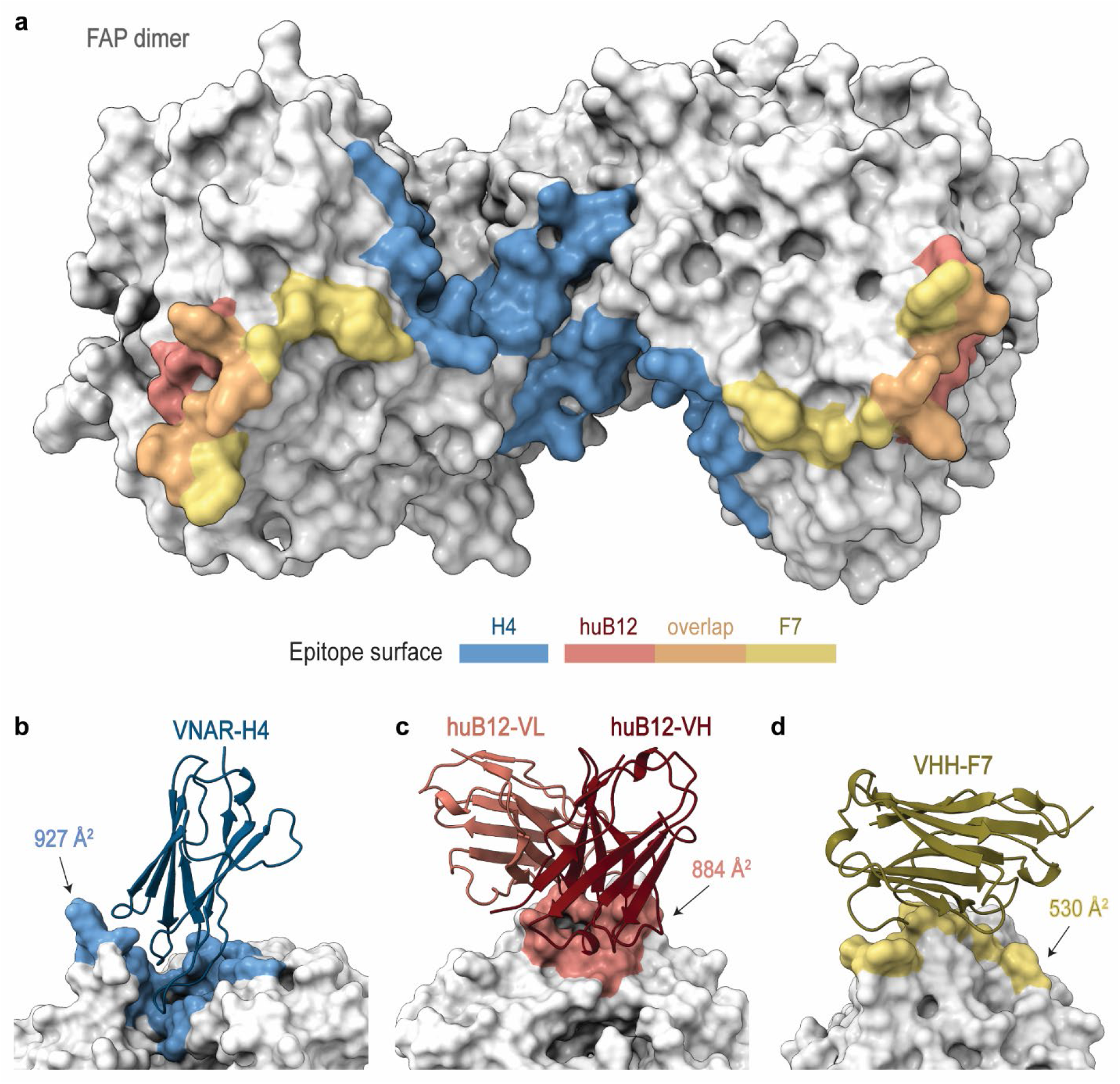
Shark VNAR-H4, humanized V_H_V_L_-huB12, and camelid VHH-F7 contact areas mapped onto FAP. (**a**) Cryo-EM surface representations of epitope mapping of VNAR-H4 (blue), huB12 (red), and VHH-F7 (yellow) along with the epitope overlap of huB12 and F7 (orange) all onto FAP homodimer (grey). Close-up ribbon representations of H4 (**b**); hub12 heavy chain (V_H_, dark red) and light chain (V_L_, light red) (**c**); and F7 (**d**) with corresponding epitope buried surface areas (BSAs).

### VNAR-H4 binds the dimer interface through a network of hydrogen bonds

Despite its compact size, the 12kDa VNAR H4 exhibited the largest BSA. Our cryo-EM reconstruction shows that the VNAR inserts into the interprotomer cleft through an extended paratope dominated by the CDR3 loop (**Fig. 2a-b**), which is consistent with previously described non-canonical VNAR engagement strategies that enable access to complex epitopes.^16,17,25,26^ Our prior X-ray crystallographic analyses indicate that VNARs engage antigens either through a conventional antibody-like mechanism with a reduced footprint or through a distinct VNAR-specific binding mode.^17,26^ H4 exemplifies the latter by inserting into the FAP dimer cleft while forming an extensive interaction interface.

**Figure 2.**
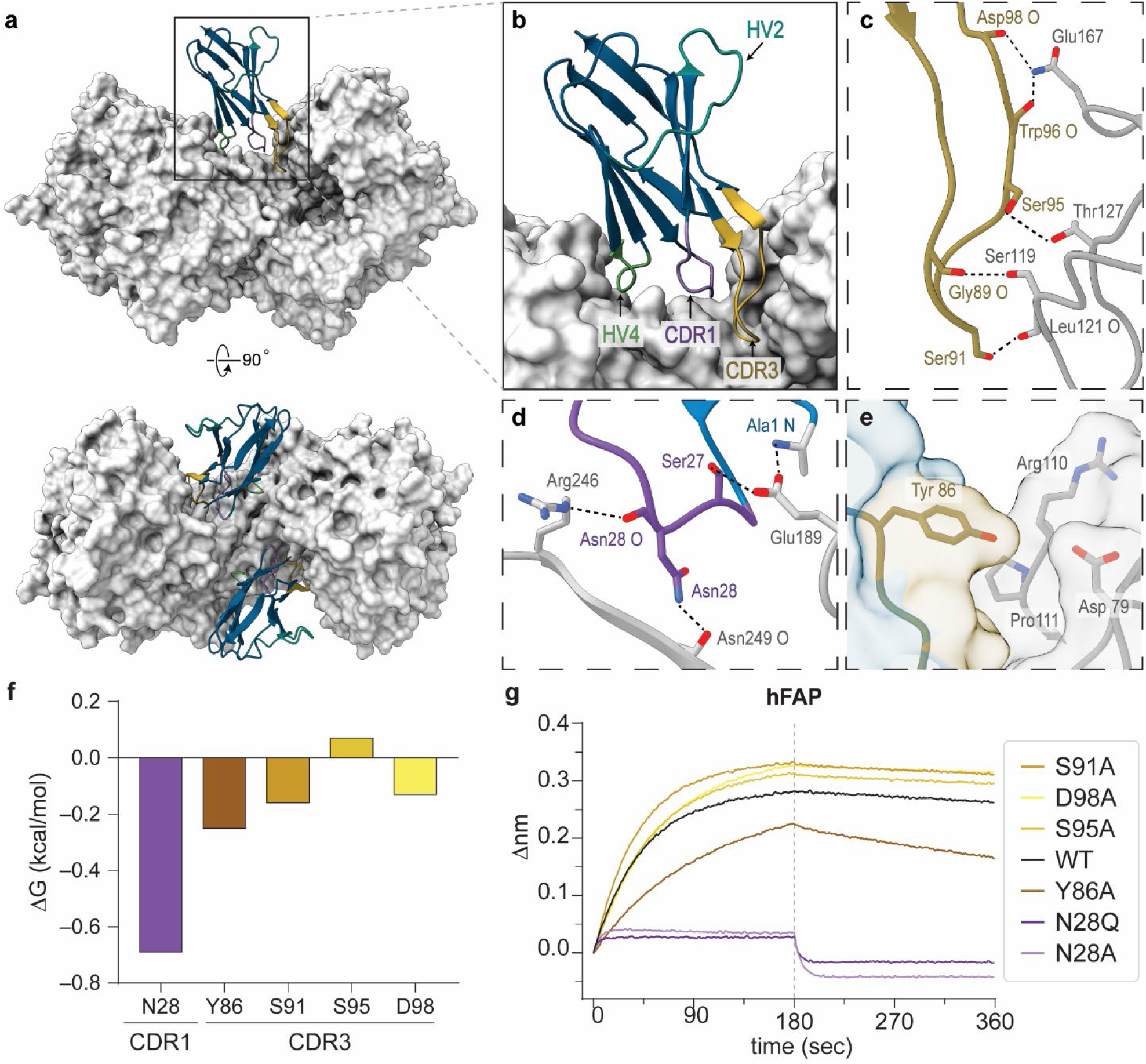
VNAR-H4 recognition is driven by CDR polar contacts. (**a**) Front and top views of the FAP–H4 complex with FAP rendered as a surface (grey). (**b**) A zoom-in of the interface highlights CDR3 (yellow), CDR1 (purple), HV4 (green), and HV2 (cyan). (**c**) The CDR3 loop contains a serine-rich region forming multiple polar contacts. (**d**) The CDR1 forms multiple Asn28 interactions with FAP. (**e**) Tyr86 buries surrounding surface area (38 Å²). (**f**) Estimated ΔG values (kcal/mol) for residues selected for cross-mutagenesis (PISA). (**g**) Real-time association and dissociation sensorgrams for H4-Fc WT and point-mutant variants (250 nM) binding to human FAP (FAP) with reports for mouse FAP (mFAP) shown in **Supp.** Fig. 2.

The FAP–H4 structure reveals an extensive hydrogen bonding network along the dimer interface (**Fig. 2c–d**), with numerous polar contacts contributed by the serine-rich CDR3 loop. In particular, CDR3 residues Gly89, Ser91, Ser95, Trp96, and Asp98 form hydrogen bonds with FAP backbone and side-chain atoms, including Ser119, Leu121, and Gln167 (**Fig. 2c**). CDR1 further reinforces the interaction through Asn28, which forms hydrogen bonds with the backbone carbonyl of Asn249 and the side chain of Arg246 on FAP. Consequently, the hydrogen bonding network of Asn28 represents the most energetically favorable contact identified by PISA (ΔG = −0.69 kcal mol^⁻^¹) (**Fig. 2d,f**).^27^ These interactions are further supported by a salt bridge between the positively charged H4 N-terminal Ala1 amine and Glu189 side chain of FAP (**Fig. 2d**). The CDR architecture is constrained by the conserved Cys29–Cys93 disulfide linking CDR1 and CDR3,^19^ together with the intradomain Cys22–Cys83 disulfide within the VNAR β-barrel core, maintaining a compact fold compatible with insertion into the dimer groove. The HV loops contribute minimally to FAP engagement. HV4 forms a single polar contact between Ser61 and Glu242, whereas HV2 is distal to the interface and makes no direct contact with FAP (**Fig. 2b**).

Using our atomic model, we identified key residues involved in H4-FAP engagement based on notable ΔG and BSA contributions (**Fig. 2e-f**). We selected Asn28, Tyr86, Ser91, and Asp98 for site-specific mutagenesis followed by BLI to assess their contributions to binding. Even though most interactions originate from the CDR3, mutating Asn28 in CDR1 to alanine (N28A) completely abrogated binding (**Fig. 2g**). Interestingly, a conservative mutation to glutamine (N28Q) also eliminated binding. These observations support that Asn28 is critical for electrostatic and energetic effects as observed in our atomic model (**Fig. 2d**). Mutating Tyr86 (Y86A) substantially reduced FAP binding (**Fig. 2 g**), consistent with its significant BSA contribution (∼38 Å²) and role in stabilizing local hydrophobic packing at the FAP interface (**Fig. 2e**). In contrast, alanine mutations at Ser91, Ser95, and Asp98 produced only slight effects (**Fig. 2g**), suggesting these positions are not major contributors for FAP recognition. Additional analysis of mFAP further supported Asn28 as essential, but additional species-dependent stabilizing effects from Tyr86, Ser91, and Ser95, highlighting subtle differences in the local environment of the epitope (**Supplementary Fig. 2**).

### Structural differences in DPP4 dictate H4 specificity

Our previous specificity experiments found that H4-Fc did not recognize recombinant DPP4 immobilized on a BLI sensor or cell surface DPP4 by flow cytometry.^19^ To investigate the structural basis of the specificity of H4 for FAP, we mapped the H4 epitope onto a structural alignment of FAP and DPP4 (PDB: 1NU6) (**Supplementary Fig. 3A**).^28^ Although the overall architecture is conserved, multiple DPP4 substitutions cluster at the H4 CDR3 interface which we predict to alter both steric accessibility and local interaction geometry. Most prominently, FAP Arg148 is replaced by a glycosylated Asn150 in DPP4, introducing an N-linked glycan likely to sterically hinder CDR3 engagement, including the chemically relevant Trp96 and Tyr86. Furthermore, FAP Pro149 to DPP4 Asn159 may alter the local environment supporting aromatic stabilization involving H4 Tyr86 in CDR3, and the substitution of FAP Gln167, which forms a hydrogen bond to Trp96 carbonyl group, while DPP4 Asn169 may further shift side-chain positioning and contact geometry at this site (**Supplementary Fig. 3B**). Additional substitutions occur near the Asn28 interaction site, including FAP Arg246, Ile248, and Asn249 corresponding to DPP4 Lys250, Val252, and Arg253, respectively, which could collectively alter the local electrostatic environment and packing around the interface (**Supplementary Fig. 3C**). Further changes are predicted to weaken specific polar contacts observed in the FAP–H4 structure, including FAP Ser119 to DPP4 Val121, which would remove the backbone-mediated interaction with CDR3 Gly89 backbone carbonyl (ΔG = -.18 kcal mol^⁻^¹).While some interactions appear chemically plausible in the DPP4 model, the combined effects of glycan occlusion, altered side-chain chemistry, and local geometric differences are predicted to disrupt the interaction network observed in the FAP–H4 complex, providing a structural rationale for H4 specificity toward FAP over DPP4.

### HuB12 and F7 converge on a shared dominant epitope

Cryo-EM reconstructions of the FAP complexes with huB12 and F7 revealed that both antibodies bind an exposed region of the FAP β-propeller domain distinct from the recessed dimer interface targeted by H4 (**Fig. 1**). Despite arising from different scaffolds, huB12 and F7 engage overlapping surfaces at the outer edge of the β-propeller domain, indicating convergent recognition of a dominant accessible epitope. The huB12 Fab forms a broad, predominantly aromatic interface that buries approximately 884 Å² of solvent-accessible surface area on FAP as calculated by PISA. Interaction analysis reveals extensive π-mediated contacts contributed by tyrosine- and phenylalanine-rich CDR residues (**Fig. 3a-b)**. Notably, heavy chain residues Phe27, Phe29, Tyr32, Tyr100, Phe101, and Tyr103, together with light chain Tyr32, Tyr49, Tyr91, and Trp96, form an aromatic packing network with FAP residues Tyr271 and Tyr274 (**Fig. 3c-d**). In addition to these stacking interactions, huB12 forms multiple hydrogen bonds across the interface. Heavy chain CDR3 residues Tyr94, Asp99, Tyr100, Asp102, and Tyr103, together with light chain Tyr32 and Tyr91 and heavy chain Arg46 and Ser31, contribute to a polar interaction network, while CDR2 residues Asn52 and Ser53 form additional contacts with FAP Glu302 (**Fig. 3c-e**). The relatively planar topology of this interface supports strong surface complementarity and π–π stabilization.

**Figure 3.**
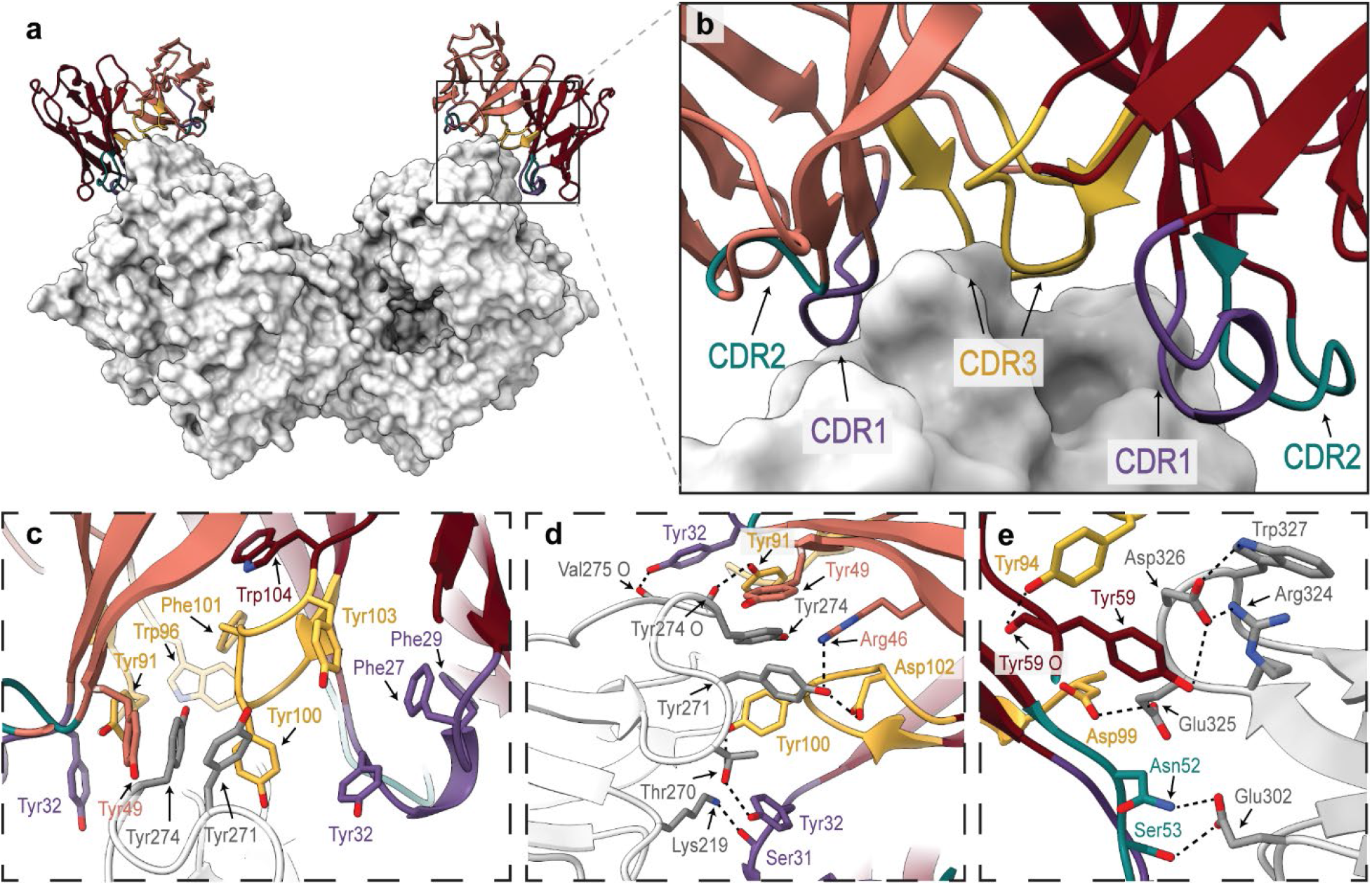
HuB12 forms aromatic-rich interaction network along β-propeller blade. (**a**) Front view of the FAP–huB12 complex is shown in. (**b**) A close-up view of the huB12 recognition site highlights CDR1 (purple), CDR2 (dark green), and CDR3 (yellow) for V_H_ (red) and V_L_ (pink). (**c**) Close-up of the aromatic interface formed huB12 CDR residues packing against FAP residues. (**d**) Additional aromatic and polar interactions formed by huB12 heavy- and light-chain residues across the FAP surface. (**e**) Polar interaction network involving huB12 CDR residues and FAP acidic/basic residues near the edge of the β-propeller epitope.

The VHH F7 engages a similar epitope with a smaller BSA of approximately 530 Å² (**Fig. 4a**). Structural analysis supports a compact interaction network mediated primarily by the CDR loops (**Fig. 4b**). Within CDR1, Tyr59 contributes aromatic packing and π–π stacking with FAP Trp327 (**Fig. 4c**). Additional aromatic residues in CDR2, including Tyr35 and Trp33, are oriented toward the interface and may support local packing interactions (**Fig. 4c**). In CDR3, Tyr110 is positioned to form polar and aromatic contacts with FAP Tyr274 and Tyr271, while backbone carbonyl and side-chain groups from Pro103, Val104, Gly105, Asn106, and Gly107 form hydrogen bonds with FAP residues Asp376, Gln328, Tyr274, and Tyr271, respectively (**Fig. 4d**). Overall, the F7 paratope achieves tight packing against the exposed β-propeller surface through a reduced but similarly aromatic interaction strategy.

**Figure 4.**
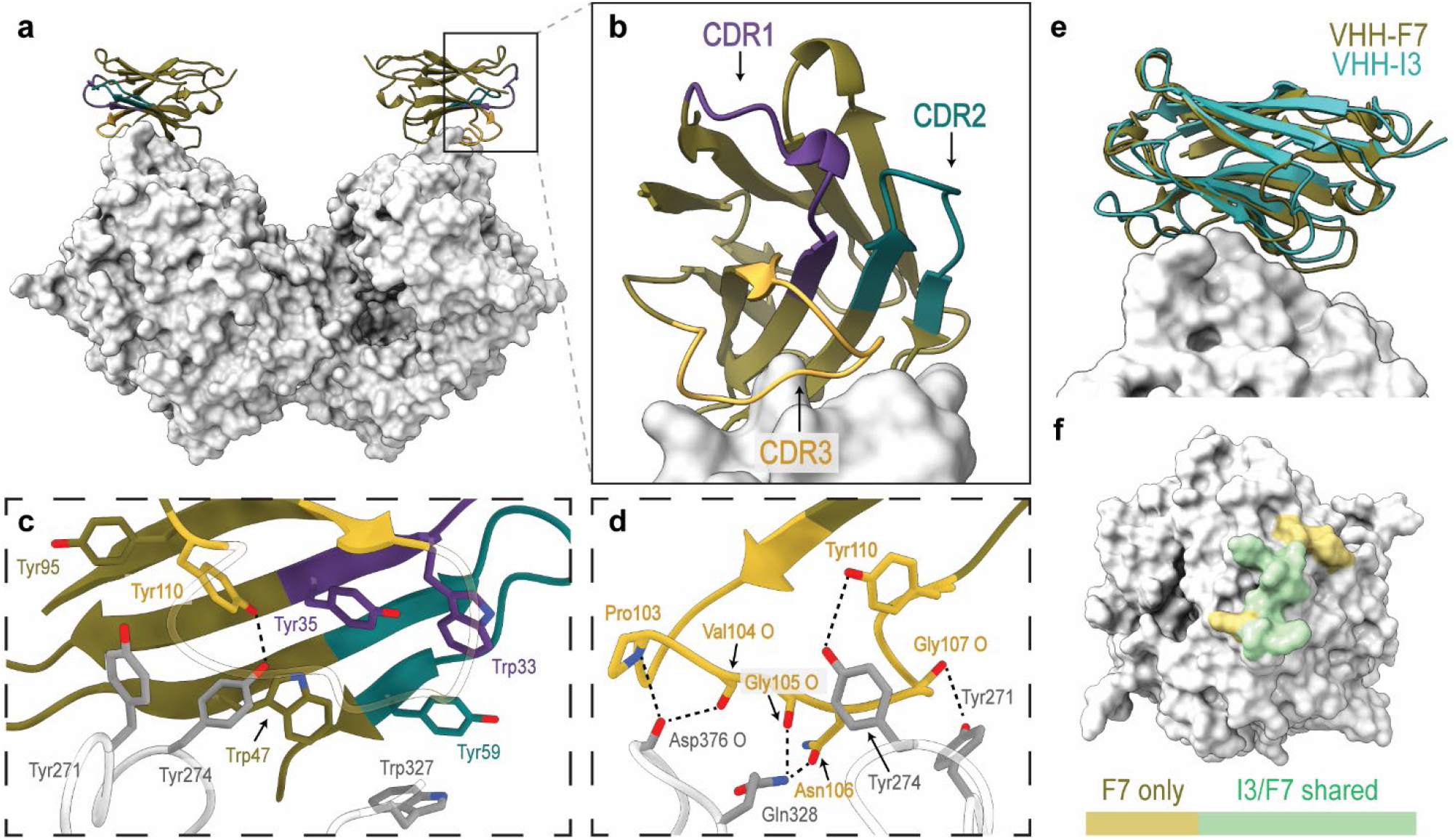
F7 engages the shared FAP β-propeller epitope through a compact aromatic interface. (**a**) Front view of the FAP–F7 complex. (**b**) A close-up view of the F7 recognition site highlights CDR1 (purple), CDR2 (green), and CDR3 (yellow). Close-up of aromatic residues at the F7-FAP interface (**c**) and of the F7 CDR3 interface (**d**) show polar contacts between F7 and FAP residues along with potential π-π stacking. (**e**) Structural alignment of FAP-bound F7 and I3 (PDB: 9DVR, cyan) reveals similar binding orientations at the β-propeller surface. (**f**) Surface mapping shows that the I3 epitope is contained within the broader F7 epitope. Residues contacted only by F7 are shown in yellow, and residues shared by F7 and I3 are shown in green.

Although the modeled F7 interface has a more favorable predicted interaction energy per domain (−6.8 kcal mol^⁻^¹) than either the huB12 V_H_ or V_L_ interface domains (−3.9 and −4.0 kcal mol^⁻^¹, respectively), huB12 distributes its contacts across both variable domains and forms a substantially larger composite interface. Nevertheless, F7-Fc binds FAP with higher apparent affinity than huB12 despite its smaller modeled contact area. This difference may reflect particularly favorable local interactions, effects of construct valency, or contacts not resolved in the primary VHH–FAP model. In support of this possibility, a subset of particles exhibits additional density adjacent to the complex (**Supplementary Fig. 4A-B**), consistent with a secondary interaction that is not captured in the primary epitope model. The size and shape of this density are compatible with an Fc domain, and preliminary fitting places the Fc near the FAP surface, although the local resolution is insufficient to confirm binding orientation and contribution (**Supplementary Fig. 4C**).

Both huB12 and F7 engage a solvent-exposed region of FAP that has been identified in a prior structural study. Structural alignment with the previously reported camelid antibody I3 (PDB: 9DVR),^29^ which was also identified from a naïve VHH phage display library, confirms a highly similar binding orientation with significant overlap (**Fig. 4e-f**, **Supplementary Fig. 5**). As with huB12 and F7, FAP residues Tyr271 and Tyr274 contribute to the ability of I3 to engage FAP. Overall, these structures indicate that humanized Fab and camelid VHH scaffolds independently converge on a shared peripheral FAP epitope dominated by aromatic and polar contacts, which contrasts the recessed binding mode of the shark VNAR H4.

### FAP undergoes local structural changes

Upon F7 binding, FAP Trp327 adopts a rotamer conformation that packs into an aromatic-rich pocket formed principally by F7 residues Trp47 and Pro103, with Tyr59 contributing to the surrounding hydrophobic environment (**Fig. 4c**). These contacts likely stabilize the outward-facing conformation of Trp327 relative to the FAP-H4 and FAP-huB12 structures (**Supplemental Fig. 6**). Conversely, huB12-FAP Trp327 rotates inward, likely due to steric clashes upon VH binding. This rotation is accompanied by a coupled change in the neighboring Arg324 rotamer, which would otherwise sterically clash with the huB12-FAP Trp327 indole ring. Other conformational changes include a slight 2.22-degree rotation consistent with the FAP dimer “opening” in the H4-bound complex, relative to the other FAP structures. This subtle rotation is likely a result of two H4 domains binding to the dimer interface shifting both FAP β-propeller domains outward.

The comparison of the complexed atomic models to the apo-FAP crystal structure (PDB: 1Z68) reveals a noticeable difference at Tyr432-Pro433, which sits within a protruding peripheral loop along the lateral surface of the outermost β-propeller blade (**Supplemental Fig. 7A**).^7^ The local backbone geometry further differs here as the Tyr432-Pro433 peptide bond adopts a highly twisted geometry in the apo-FAP crystal structure, which contrasts the cis-conformation in all our antibody-bound FAP structures (**Supplemental Fig. 7B**). The altered position of this loop and peptide in the apo-FAP structure may reflect crystal packing contacts as the solvent-exposed residues exhibit stabilizing interactions between neighboring molecules of a symmetrized FAP-forming lattice (**Supplemental Fig. 7C-D**).

### Additional immune-derived VNARs recognize FAP surfaces outside the shared β-propeller epitope

We next analyzed three FAP-VNAR complexes (H4-Fc, H15-Fc, and NGS2405-Fc) by hydroxyl radical foot printing MS to investigate if our other VNARs identified from the immunized library recognized the shared β-propeller epitope. The data generated is akin to hydrogen-deuterium exchange mass spectrometry, which reports on solvent accessibility and epitope engagement in solution. Comparison with the cryo-EM structure showed that the H4 footprint broadly overlaps the experimentally determined epitope, although the footprinting-derived docking model did not accurately reproduce the VNAR orientation or residue-level interactions. Despite this difference in precision, the results of the FAP-H4 complex support the epitope mapping data from our structure and thus gave us the foundation to believe the other VNAR binding regions determined by HRF-MS were valid predictions (**Fig. 5a**). Both VNARs recognize epitopes distant from the dimer interface and the shared β-propeller region. For NGS2405, the epitope is located at the N-terminus close to plasma membrane while the epitope for H15 is near the C-terminus on the α/β-hydrolase domain. Our binning studies support that H17 and sibrotuzumab epitopes, while not identical, are in the vicinity of H15. These findings suggest that immune-derived antibodies, including shark-derived VNARs and sibrotuzumab, appear to access alternative, structurally diverse epitopes not targeted by the naïve phage display-derived antibodies examined here (**Fig. 5b-c**).

**Figure 5.**
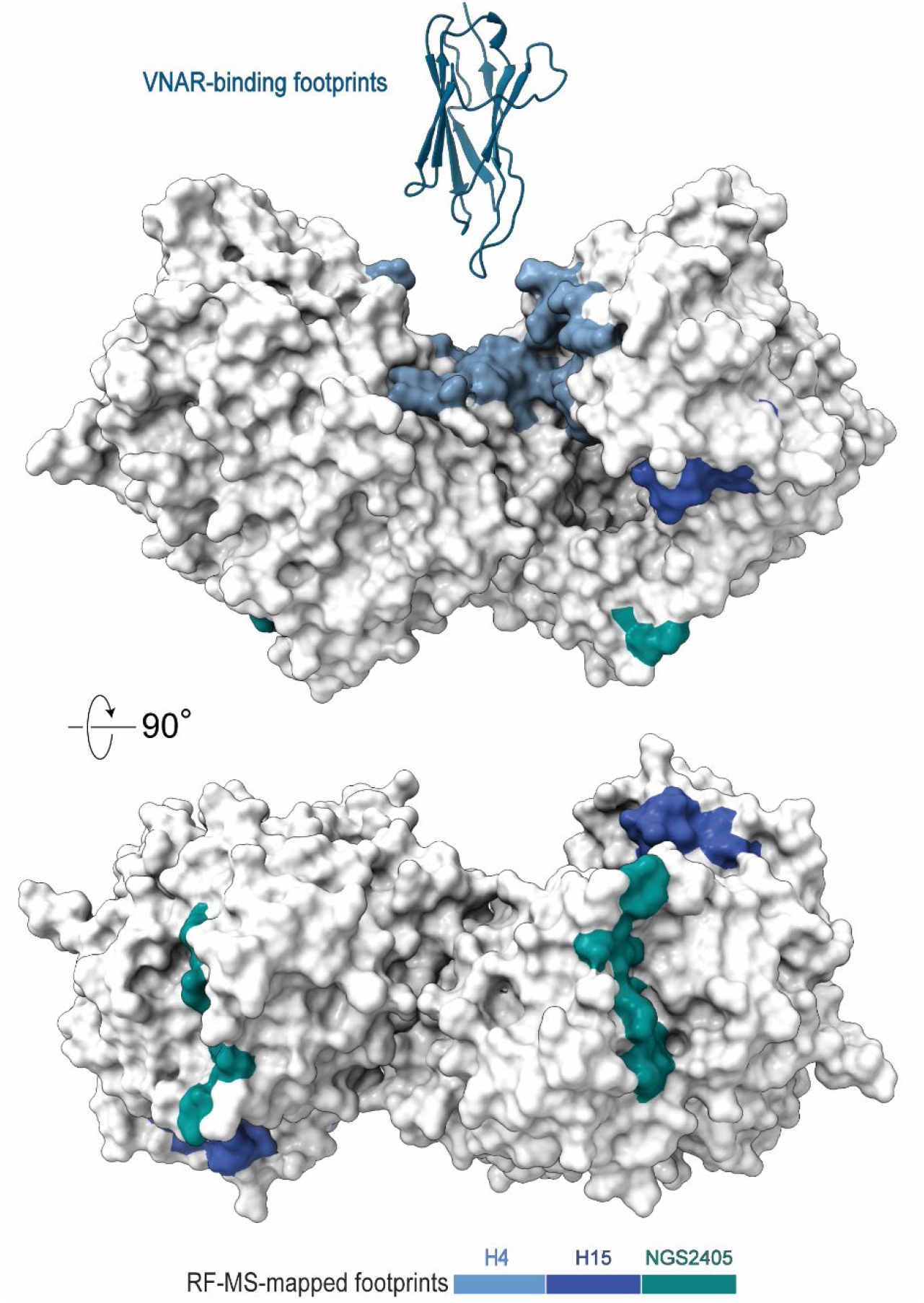
Radical footprint mass spectrometry reveals additional VNAR binding epitopes on FAP. Surface-mapped footprinting data are shown on FAP with the H4 cryo-EM epitope highlighted in blue. Regions protected in radical footprinting experiments corresponding to H15 (indigo) and NGS2405 (teal) VNARs are also shown, revealing additional solvent-exposed binding surfaces of FAP. Highlighted residues indicate regions exhibiting significant protection or deprotection upon VNAR binding and define probable binding surfaces but do not establish the orientation or atomic contacts of the bound VNAR.

### Mapped VNAR epitopes occur within conserved regions of FAP

To investigate if VNARs prefer conserved domains, we curated FAP protein sequences from 41 chordates spanning multiple classes and performed a sequence alignment which allowed us to develop a conservation map of FAP that was applied to our structure (**Fig. 6**). Surprisingly, FAP was highly homologous across all the species analyzed (**Supplemental Table 2**). The residues at the dimer interface and the epitope of H4 were found to be nearly identical across all species with few point mutations. Glu242 of FAP which forms a hydrogen bond with Ser61 of H4 was replaced with a Gly residue in rodents, which potentially explains the differences in affinity of H4 for human and mouse FAP (**Fig. 6b**). The H15 epitope was completely conserved among primates and moderately conserved across mammals. In mice, there were radical substitutions where Ala72 and Asn74 in humans were replaced by a Glu and Asp, respectively. These substitutions introduce negatively charged residues, which likely account for the lack of cross-reactivity of H15, H17, and sibrotuzumab with murine FAP. For NGS2405, some variability was observed, however, the complete epitope was conserved in nearly all the mammals with several variants in the rodents (**Supplemental Fig. 8**). Unlike the VNAR epitopes examined here, the shared huB12, F7, and I3 epitope exhibited greater sequence variability across species and mammals (**Fig 6c-d**). Human FAP Tyr274, which contributes to the huB12, F7, and I3 interfaces,^29^ is replaced by histidine in rodents. However, huB12 retains cross-reactivity with murine FAP whereas F7 does not, demonstrating that this substitution alone is insufficient to explain the observed species specificity. These findings overall indicate that VNARs preferentially recognize highly conserved regions of FAP.

**Figure 6.**
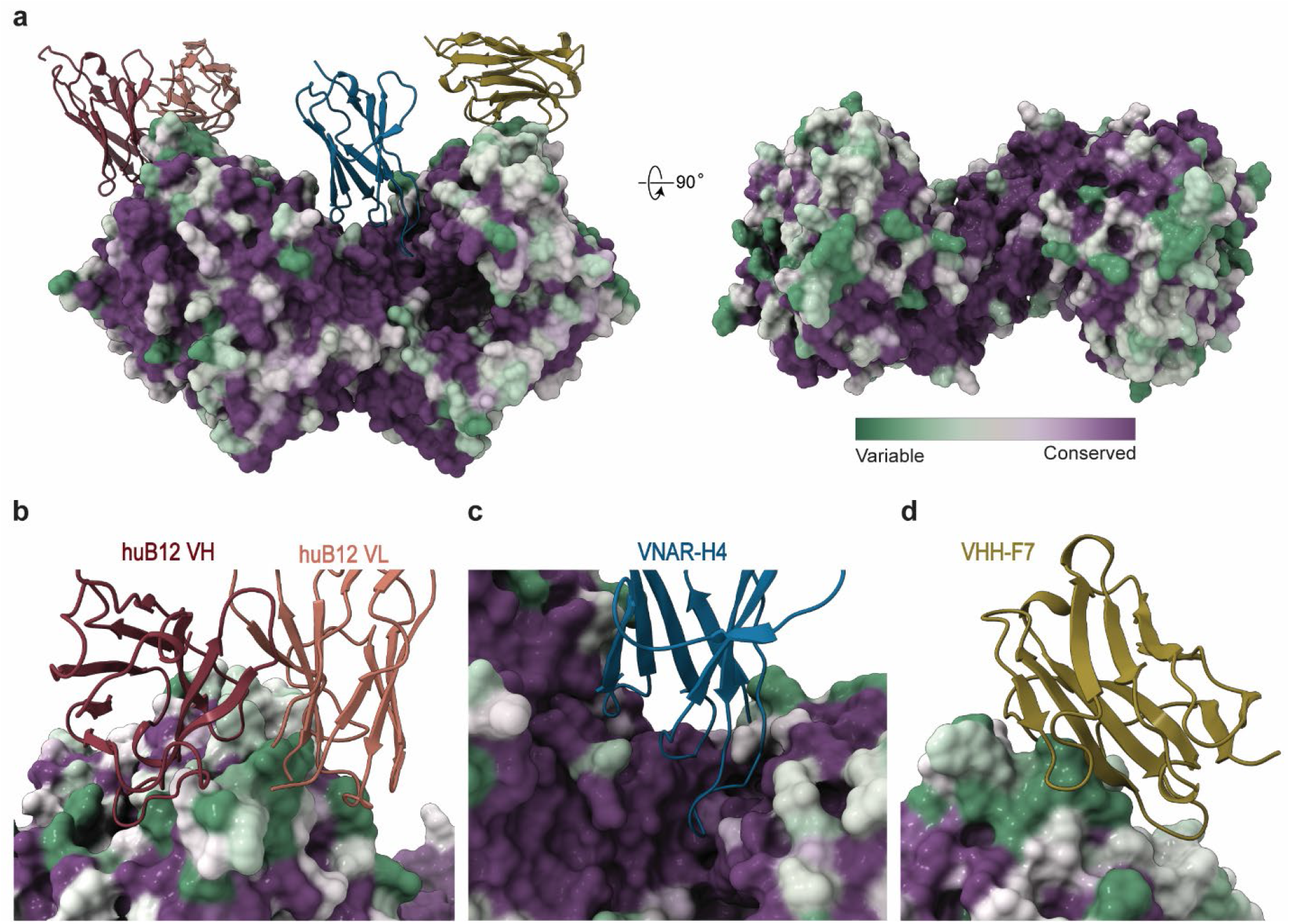
Conservation of FAP epitopes. (**a**) FAP surface conservation mapped by ConSurf reveals distinct epitope conservation across the antibody-bound complexes with variable residues shown in green, average residues in white, and highly conserved residues in purple. Close-up display of the FAP binding sites for humanized huB12 (**b**), shark VNAR-H4 (**c**), and camelid VHH-F7 (**d**).

## Discussion

The increased expression of FAP on activated fibroblasts in the reactive stroma is a distinguishing feature of cancer, fibrosis, and chronic inflammatory conditions, making FAP an attractive biomarker and therapeutic target.^6^ Nevertheless, relatively few anti-FAP antibodies have been investigated compared with small-molecule FAPIs.^30,31^ Although several structures have characterized apo or small-molecule-bound FAP, the FAP-I3 cryo-EM study remains the only previously published structural analysis of an antibody-bound FAP complex.^7,29^ In contrast, FAP’s closest ancestral homolog DPP4 has been extensively characterized with both small molecules and antibodies, providing a structural basis for rational therapeutic design that remains largely absent for FAP.^8,9,32,33^ Here, we address this gap through a comparative structural analysis of FAP recognition by humanized, camelid, and shark-derived antibodies. To our knowledge, the FAP-H4 complex provides the first cryo-EM structure of a VNAR and reveals a mode of FAP recognition not observed for previously characterized antibody scaffolds.

Because dimerization is required for FAP activity, the dimer interface is among its most highly conserved regions, together with the active site. Several FAP residues that contribute strongly to H4 binding are conserved across species but differ in the paralog DPP4, consistent with the cross-reactivity of H4 toward murine FAP and its lack of detectable DPP4 binding. Moreover, radical footprinting MS findings indicated that the H15 and NGS2405 VNAR epitopes are more highly conserved than the huB12, F7, and I3 epitope region. Previous studies have similarly demonstrated VNAR engagement with well-conserved epitopes, including a SARS-CoV-2-receptor-binding domain retained across multiple variants.^16–18,25,30,32^ These prior results, combined with our structural comparisons, support the broader capacity of VNARs to target evolutionarily constrained and otherwise underexplored antigen surfaces. This property may be particularly relevant for targeting FAP-positive stroma, a common feature across heterogeneous tumor contexts that can actively promote therapeutic resistance.^19,34–37^ Thus, VNARs may offer a dual advantage: directing therapy toward a resistance-supporting stromal compartment while, in the case of H4, raising the barrier to mutational escape by engaging a highly conserved interface required for FAP enzymatic activity. This proposed advantage requires direct testing and would not preclude antigen loss or other resistance mechanisms.

The non-overlapping epitopes identified here, furthermore, provide a structural basis for developing biparatopic or multispecific FAP binders. Combining binders that engage spatially distinct regions of FAP could broaden recognition across cellular contexts and potentially mitigate the effects of receptor clustering or differences in FAP presentation on the cell surface. Multiepitope engagement may also confer new functional properties, as demonstrated by the distinct receptor-clustering geometry induced by biparatopic HER2 binding.^38^ Biparatopic anti-TNF-α VNARs have demonstrated that combining two epitope specificities can substantially enhance binding affinity and neutralization potency.^17^ The separation between the H4 dimer-interface epitope and the exposed sites recognized by huB12 and F7 could therefore permit simultaneous engagement of FAP through complementary binding geometries. However, the structures establish epitope separation rather than functional synergy. The distinct internalization profiles of H4, huB12, and F7 nevertheless support testing interface- and exposed-site combinations that integrate binding and trafficking, while contributions of format-dependent variables remain to be resolved.

Interestingly, huB12, F7, and I3 independently converged on the same accessible β-propeller region. All three antibodies were isolated from naïve human or camelid libraries, whereas the immune-derived VNARs H4, H15, and NGS2405 recognize distinct surfaces. The immune-derived murine precursor of sibrotuzumab, F19, similarly recognizes a region overlapping those targeted by the shark-derived H15 and H17 antibodies. This observed pattern raises the possibility that discovery strategy influences which FAP surfaces are sampled. However, the number of characterized antibodies remains limited, and scaffold type, library composition, and selection conditions differ among these examples. The apparent association could, as a result, be coincidental. Alternatively, naïve library selections may preferentially recover binders to highly accessible, selection-enriched surfaces; meanwhile, immune repertoires may sample more recessed or conformationally constrained epitopes. Parallel selection from naïve and immunized libraries under matched antigen-presentation and panning conditions could directly test this hypothesis. If this pattern proves reproducible, combining these discovery approaches could expand the range of targetable FAP epitopes available for multiepitope antibody development. Overall, these findings characterize distinct modes of FAP recognition and suggest that antibody scaffold and discovery strategy can broaden epitope access for rational design.

## Methods

### Antibody production

DNA sequences encoding anti-FAP antibodies were codon optimized for expression in Chinese hamster (*Cricetulus* griseus) systems and synthesized as double stranded DNA gBlocks (Integrated DNA Technologies, Coralville, IA), prior to integration into a mammalian expression vector encoding the Fc domain of human IgG1 (TGEX-SCblue, Antibody Design Labs) by homologous recombination, using an In-Fusion HD cloning kit (Takara Bio USA, San Jose, CA). Plasmid constructs encoding antibodies were transfected into ExpiCHO-S cells using an Expifectamine CHO Transfection Kit (Gibco, Thermo Fisher Scientific, Waltham, MA). Transfection and ExpiCHO-S cell culture maintenance was performed as recommended by the manufacturer. 12-14 days post transfection, cultures were harvested and centrifuged at 2000xRCF for 10 min at 4 °C. Protein-containing supernatant was collected and clarified by centrifugation at 20,000xRCF for 30 min at 4 °C, supplemented with 300mM NaCl and 20mM Na_2_PO_4,_ pH was adjusted to 6.8, and solutions were passed through a 0.45 µm sterile filter. Clarified ExpiCHO-S supernatant was run through PBS equilibrated MabSelect PrismA columns (Cytiva, Marlborough, MA) at 1 column volume (CV) per min. Unbound protein was washed from the column using 10 CVs of PBS. Antibodies bound to the column were eluted in 8 CVs of 100 mM glycine at pH 3.0. Eluate was immediately neutralized using 2 M Tris HCl at pH 8.6. Eluates were concentrated and buffer exchanged into PBS using an Amicon stirred chamber with a 30 kDa MWCO ultrafiltration membrane (Millipore). Size exclusion chromatography was performed on an ÄKTApure fast protein liquid chromatography system using a HiLoad 16/600 Superdex 200 pg (Cytiva) column equilibrated with PBS. Samples were loaded and fractionated using a mobile phase of PBS at 0.5 mL/min; chromatograms were obtained by monitoring UV absorbance at 280 nm. Eluted protein fractions contributing to a single peak of UV absorbance corresponding to the theoretical molecular mass of each antibody were collected and pooled. Eluates were aliquoted, flash frozen and stored at -80°C.

### Biolayer Interferometry

BLI studies were performed in an assay buffer consisting of PBS supplemented with 1% BSA. Recombinant human FAP (FAP-H82Q56, Acro Biosystems) and recombinant mFAP (FAP-M82Q8, Acro Biosystems) were purchased pre-biotinylated. For determination of purified antibody dissociation constants, hydrated SAX sensors were equilibrated for 30 secs in assay buffer before loading of biotinylated bait protein (30 nM) for 90 sec, followed by a baseline equilibration for 30 sec in assay buffer. Association of antibodies (250 nM) to biosensors with immobilized bait protein was monitored for 3 minutes, dissociation in assay buffer was monitored for an equivalent period of time. Each experiment included a reference well with no analyte, to ensure specificity of signal on the bait-loaded sensors. Binding affinities were determined by kinetic analysis of binding curves in the Octet Data Analysis software (v12.0.2.3), using conditions reflecting bivalent analyte and background subtracting data from reference wells. For binning assays, hydrated SAX biosensors were monitored during a 30 s baseline in assay buffer, loaded with biotinylated human FAP (30 nM, 90 s), followed by exposure to a saturating concentration of a single primary antibody (1 µM). Biosensors were then exposed to a second saturating concentration of each individual antibody (1 µM) as to determine additional association to the bait protein.

### Antibody-FAP Complex Formation

Antibody-FAP complex formation was performed at a 1:3 molar ratio using 200 μL of FAP (7 μM) and 200 μL of antibody (21 μM) in PBS buffer, with the final volume adjusted to 500 μL. The Antibody-FAP solution was incubated at room temperature for 2 hours to allow complex formation. Size exclusion chromatography was then performed, as described above, to separate the complexed and non-complexed proteins (**Supplemental Fig. 9**). The complexed peak, as determined by its elution time, was pooled and concentrated using an Amicon a 30 kDa MWCO ultrafiltration membrane (Millipore). To confirm complex formation, a 2 μg sample was run on a Bolt™ Bis-Tris PLUS gel (Invitrogen, Thermo Fisher Scientific) under reduced and non-reduced conditions (150V for 45 min). The gel was stained with GelCode™ Blue Safe Protein stain (Thermo Fisher Scientific) for 30 minutes, followed by overnight background clearing in water.

### Cryo-EM sample preparation

Purified FAP in complex with H4-Fc, huB12, and F7-Fc were diluted in PBS to final concentrations of 2.4 mg/mL, 0.6 mg/mL, and 1.3 mg/mL, respectively. Initial screening of these complexes indicated that standard ice thicknesses were insufficient for optimal particle distribution. To address this, we utilized UltrAuFoil R 1.2/1.3 holey gold support films (Quantifoil, Micro Tools GmbH). Grids were glow-discharged for 20 s at 20 mA in ambient air using a PELCO easiGlow system (Ted Pella, Inc.). A 3 uL aliquot of the sample was applied to the grid, blotted for 4 s with a blot force of +22, and vitrified in liquid ethane using a Vitrobot Mark IV (Thermo Fisher Scientific) operated at 4 °C and 100% humidity with a 0.5 s drain time. While initial trials using C-Flat holey carbon films (Protochips) improved particle visibility, 2D classification revealed significant preferred orientation, limiting reconstructions to ∼4.3 Å. Consequently, we reverted to UltrAuFoil grids and increased sample concentrations to promote thicker ice, selecting holes with prominent ice rings. This approach reduced beam-induced motion and provided a more diverse range of particle orientations (**Supplemental. Figs 10-11**), which proved essential for high-resolution reconstruction on gold supports.^39^

### Data Acquisition and analysis

Grids were screened on 200 kV Titan Arctica and Glacios cryo-TEMs (Thermo Fisher Scientific). Final high-resolution datasets were acquired on a 300 kV Titan Krios G3i equipped with a Gatan K3 direct electron detector and a BioQuantum energy filter (20 eV slit width). Data were collected in CDS-counting mode using SerialEM with a multi-shot strategy.^40^ Images were recorded at a calibrated pixel size of 0.834 Å, with a total electron dose of 80 e^⁻^Å^⁻^² fractionated over 80 frames. Nominal defocus values ranged from -0.5 to -1.5 µm. To overcome remaining orientation bias, the H4-Fc and huB12 datasets were collected with a 20° stage tilt, whereas the F7-Fc dataset was collected at 0° tilt.^41^ All movie frames were aligned, dose-weighted, and resampled to a pixel size of 1.0 Å during preprocessing. High-resolution reconstructions were performed entirely within the cisTEM software package (**Supplemental. Figs 10-12**).^42^

### Model Fitting and Validation

Initial atomic models for the FAP-antibody complexes were generated using ModelAngelo with backbone adjustments guided by the FAP crystal structure (PDB: 1Z68) and docked into the cryo-EM density.^7,43^ For the F7-Fc complex, the Fab region was modeled using an AlphaFold-guided backbone fitting to account for lower local resolution resulting from conformational flexibility in the Fc domain.^44^ Models were iteratively improved through manual building in Coot and automated real-space refinement in Phenix (**Supplemental. Figs 10-12**).^45,46^ Final model quality was assessed using MolProbity to ensure optimal Ramachandran statistics and minimal steric clashes.^47^ Protein-protein interfaces and buried surface areas (BSAs) were Protein Interfaces, Surfaces, and Assemblies (PISA) server and ChimeraX.^27,48^ Hydrogen bonds were identified based on a distance-based cutoff of 3.5 Å between donor and acceptor atoms. All structural figures and interface visualizations were rendered in ChimeraX.^48^

### Radical footprint mass spectrometry

Radical footprint mass spectrometry was performed by Immuto Scientific (Madison, WI) as a fee-for-service. Data reports are available on request.

### Conservation Analysis of Fibroblast Activation Protein FAP

FAP protein sequences were retrieved from multiple animal species. We conducted two parallel analyses: (1) using FAP sequences from diverse animal species, and (2) using FAP sequences restricted to mammalian species only. Multiple sequence alignments (MSA) were generated using MUSCLE alignment software.^49^ The crystal structure of human FAP (PDB ID: 1Z68) was used as the reference three-dimensional structure for conservation mapping. Conservation scores calculated for the primary chain were mapped identically onto the second chain to represent the complete biological assembly. Evolutionary conservation scores were computed using ConSurf.^50^ ConSurf integrates the MSA from MUSCLE with the three-dimensional protein structure to calculate position-specific conservation scores using a maximum likelihood estimation (MLE) approach. The MLE method models amino acid substitutions as a continuous-time Markov process along a phylogenetic tree, with each position assigned an evolutionary rate parameter. Slower evolutionary rates indicate higher functional or structural constraint and thus higher conservation. Conservation scores were visualized on the FAP protein surface using ChimeraX.^48^ The conservation gradient was displayed using a color scheme ranging from green (variable) to purple (conserved).

## Data Availability

The authors declare that the data supporting the conclusions of this article are included in the main text and supplementary materials. The structural data generated in this study have been deposited in the Electron Microscopy Data Bank (EMDB) under accession codes EMD-78501 (FAP-H4 shark VNAR complex), EMD-78502 (FAP-huB12 humanized IgG complex), and EMD-78503 (FAP-F7 camelid VHH complex). Atomic models are available in the Protein Data Bank (PDB) under accession codes PDB: 37US, 37UT, and 37UU, respectively. The data underlying material for Supplemental Fig. 4 is included under EMD-78503 as an additional EM map.

## Author Contributions

R.L.B., E.W.L., and T.G. performed cryo-EM acquisition and structural determination. R.L.B analyzed structural data and prepared figures. C.A.H. contributed to atomic model building and S.C. contributed to supplemental information. K.L.O. performed mutagenesis studies and determined binding affinities for proteins of interest. K.L.O. and G.S.G. performed binning studies. J.L.W. contributed to molecular cloning of mutant constructs. K.L.O. and E.W.L. expressed and purified the proteins used. T.R. and I.M.O. developed the conservation map of FAP. A.M.L, and T.G. contributed to the conceptualization and design of the study. R.L.B., K.L.O., T.R., I.M.O., A.M.L., and T.G. wrote the manuscript, with all authors contributing to its final version.

## Acknowledgments.

Cryo-EM data were collected at the Cryo-Electron Microscopy Research Center (CEMRC) in the Department of Biochemistry at the University of Wisconsin–Madison. We thank the CEMRC staff for assistance with cryo-EM data collection. We also thank Alicia Williams for editorial assistance and Matthew Stefely for assistance with figure preparation. This work was supported by the National Institutes of Health/National Cancer Institute (NIH/NCI) grants R01 CA305895, R01 CA295576, R01 CA237272, and R01 CA233562, a Prostate Cancer Foundation Challenge Award, and Andy North and Friends (A.M.L.). R.L.B. was supported by the NIH/NIGMS Molecular Biophysics Training Program under award T32 GM130550 and by the Steenbock Predoctoral Graduate Fellowship administered by the University of Wisconsin–Madison Department of Biochemistry. The content is solely the responsibility of the authors and does not necessarily represent the official views of the NIH.

## Competing Interests

The authors declare the following competing interests: A.M.L. and Joseph P. Gallant are inventors on pending international patent application PCT/US2024/041211, with the Wisconsin Alumni Research Foundation (WARF) as applicant, covering F7 and related anti-FAP single-domain antibody formats and uses. A.M.L., G.S.G., and Joseph P. Gallant are inventors on pending international patent application PCT/US2025/059927, with WARF as applicant, covering H4 and related anti-FAP VNARs and uses. A.M.L. and Hallie M. Hintz are inventors on U.S. Patent No. 11,738,050, assigned to the Regents of the University of Minnesota, covering B12/huB12 anti-FAP antibodies and related uses. The remaining authors declare no competing interests.

