## Supplemental Information for "A shark variable new antigen receptor recognizes an occluded epitope of fibroblast activation protein"

*Corresponding Authors:

### SUPPLEMENTAL FIGURES.

#### Supplemental Figure 1.

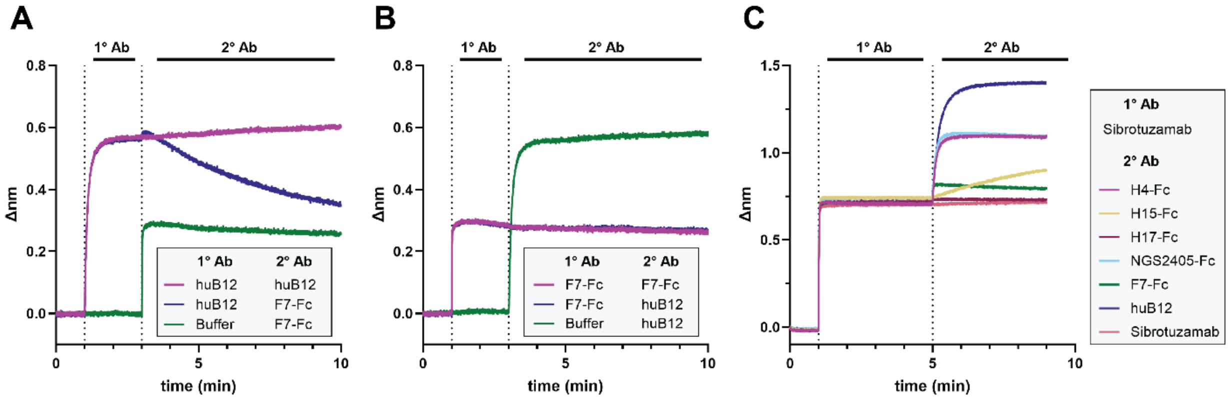


**Supplemental Figure 1. Biolayer Interferometry (BLI) epitope binning of FAP antibodies.** Binning of immobilized hFAP with (**A**) huB12, (**B**) F7-Fc, and (**C**) sibrotuzumab as the primary antibody, followed by the indicated competing antibodies.

**Supplemental Figure 2.**

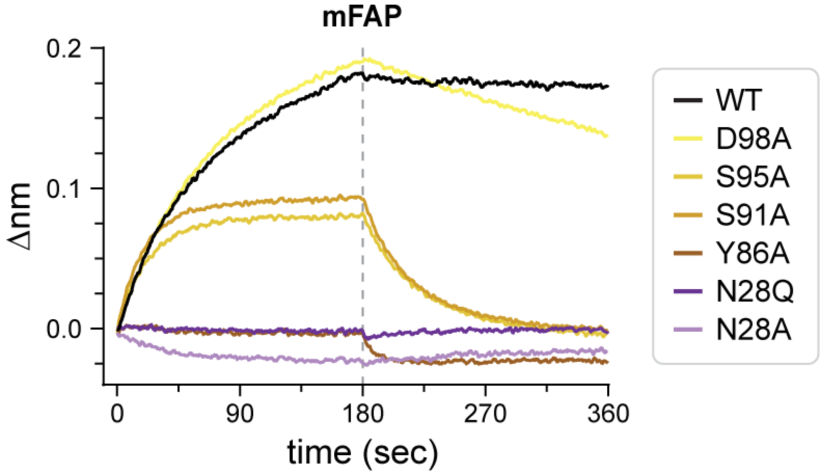


**Supplemental Figure 2. BLI affinity readings of H4 binding to mFAP.** Real-time association and dissociation sensorgrams for H4-Fc WT and point-mutant variants (250 nM) binding to mouse FAP (mFAP).

**Supplemental Figure 3.**

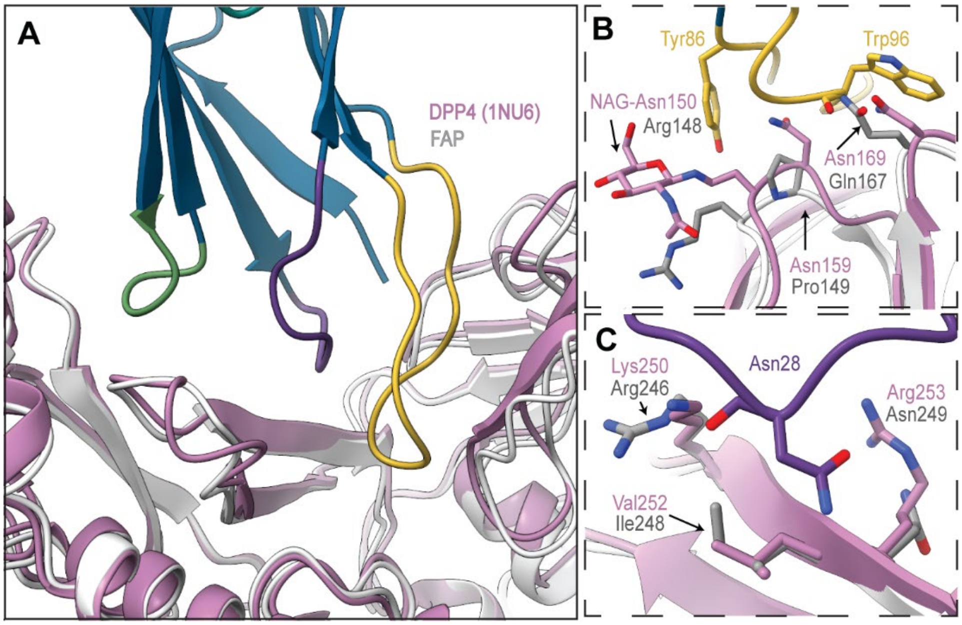


**Supplemental Figure 3. DPP4 alignment with FAP-H4.** Structural alignment of DPP4 (pink; PDB 1NU6)[1] onto the FAP–H4 complex (grayscale) (**A**) with H4 (blue) colored by CDR loop (CDR1, purple; CDR2, teal; CDR3, yellow). A close-up of the CDR3 interface (**B**) along with substitutions proximal to the CDR1-Asn28 interaction (**C**).

**Supplemental Figure 4.**

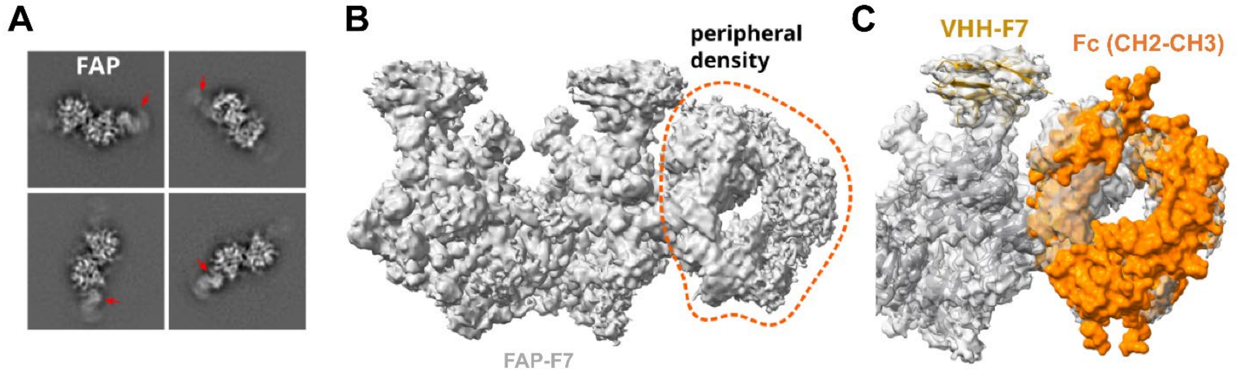


**Supplemental Figure 4. Additional EM density observed in the FAP-F7 dataset.** Two-dimensional classification of 1,520,217 picked particles identified a lower-resolution subset of 150,153 particles (9.9%) containing low-resolution density additional to the FAP-F7 data. (**A**) Representative 2D class averages showing this density adjacent to the FAP–F7 complex (red arrowheads). (**B**) Three-dimensional reconstruction of the corresponding particle subset, with the peripheral density outlined by the orange dashed contour; overall density is shown in gray. (**C**) The model fitting of fused Fc region predicted from Alpha-Fold[2] (orange) extending from the FAP-bound VHH-F7 domain (yellow).

**Supplemental Fig. 5.**

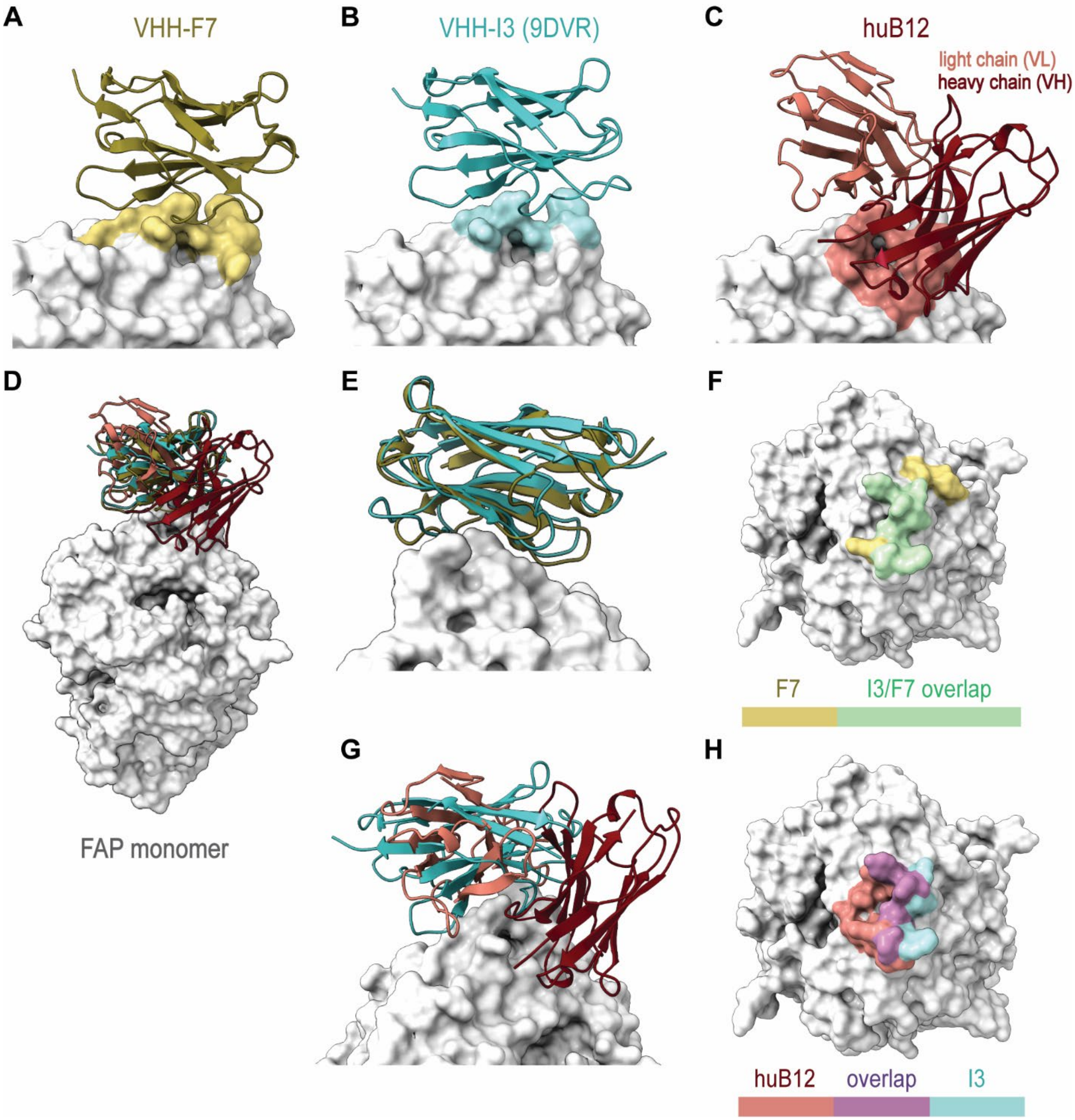


**Supplemental Fig. 5. Structural alignment of FAP-bound F7, huB12, and I3-VHH.** Binding modes for F7 (**A**), I3 (**B**, PDB: 9DVR),[3] and huB12 (**C**) mapped onto FAP monomer (grey), with corresponding epitope footprints colored on the FAP surface (yellow, cyan, red, respectively). Structural overlay of F7, I3, and huB12 on a single FAP monomer, showing convergence on the solvent-exposed β-propeller surface (**D**). Close-up alignment of F7 and I3 showing similar binding orientations (**E**) and the F7 epitope (yellow) completely overlaps the I3 epitope (green) (**F**). Overlay of F7, I3, and huB12 showing that huB12 engages the same general β-propeller region from a distinct orientation (**G**) with surface representation of the huB12 (red) and I3 (cyan) epitopes highlighting partial overlap (magenta) between the binding sites (**H**).

**Supplemental Fig. 6.**

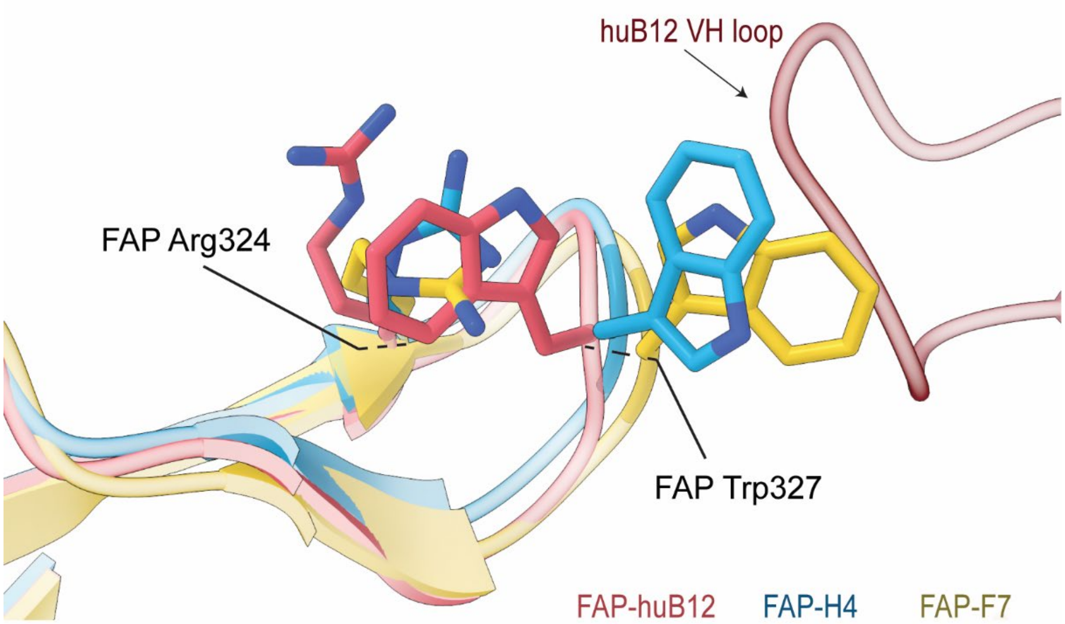


**Supplemental Fig. 6. Local rearrangements of FAP Arg324 and Trp327.** Superposition of the FAP-huB12, FAP-H4, and FAP-F7 structures showing the distinct conformations of FAP Arg324 and Trp327. FAP from the huB12, H4, and F7 complexes is shown in red, blue, and gold, respectively.

**Supplemental Fig. 7.**

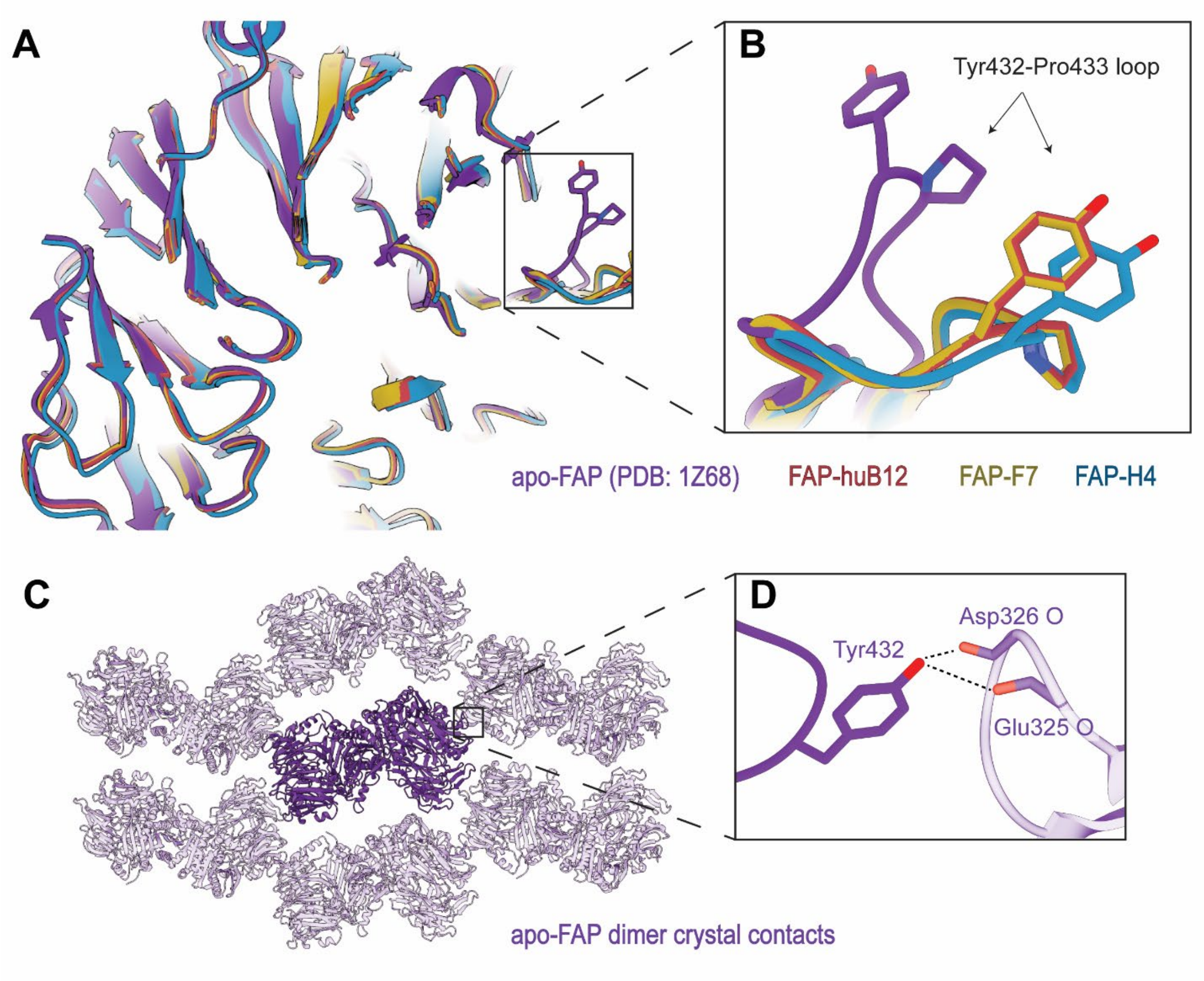


**Supplemental Fig. 7. Crystal packing may stabilize an altered Tyr432–Pro433 loop conformation in apo-FAP.** (**A**) Structural alignments of the outer β-propeller region from apo-FAP and the FAP-huB12, FAP-F7, and FAP-H4 structures. (**B**) Enlarged view of the Tyr432–Pro433 loop, highlighting its altered position in apo-FAP relative to the antibody-bound structures. Apo-FAP, FAP-huB12, FAP-F7, and FAP-H4 are shown in purple, red, gold, and blue, respectively. (**C**) Crystal-packing arrangement generated from the apo-FAP crystal structure (PDB 1Z68).[4] (**D**) Enlarged view of the crystal contact formed by Tyr432 of the focal FAP molecule and oxygen atoms (red) from Asp326 and Glu325 of a symmetry-related molecule. Dotted lines indicate intermolecular contacts within 3Å (ChimeraX).[5]

**Supplemental Figure 8.**

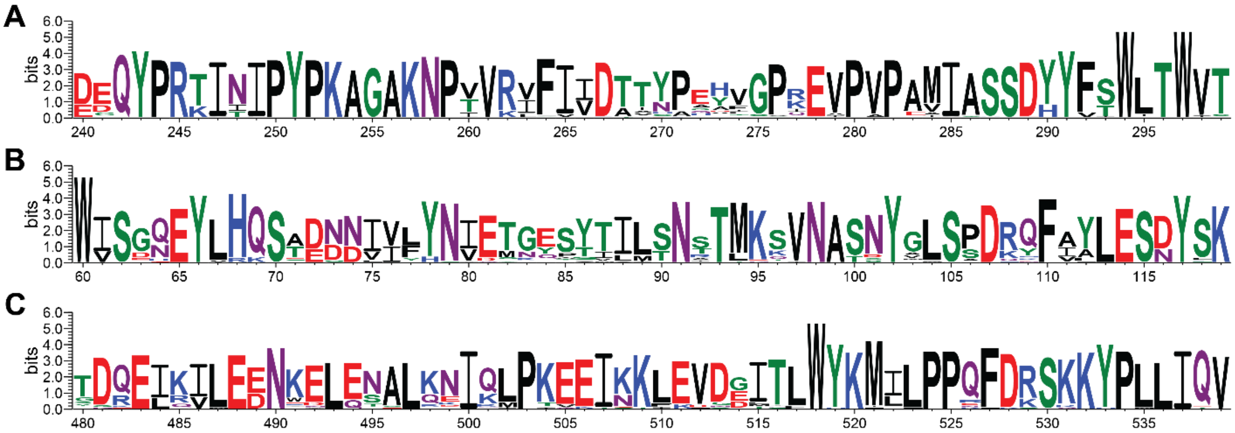


**Supplemental Figure 8. Conservation of FAP sequences within antibody epitopes across mammalian species.** Sequence logos generated from mammalian FAP amino acid sequences illustrate residue conservation within the mapped epitope regions of anti-FAP antibodies, **(A)** huB12, F7, and H4, **(B)** H15, and **(C)** NGS2405.

**Supplemental Fig. 9.**

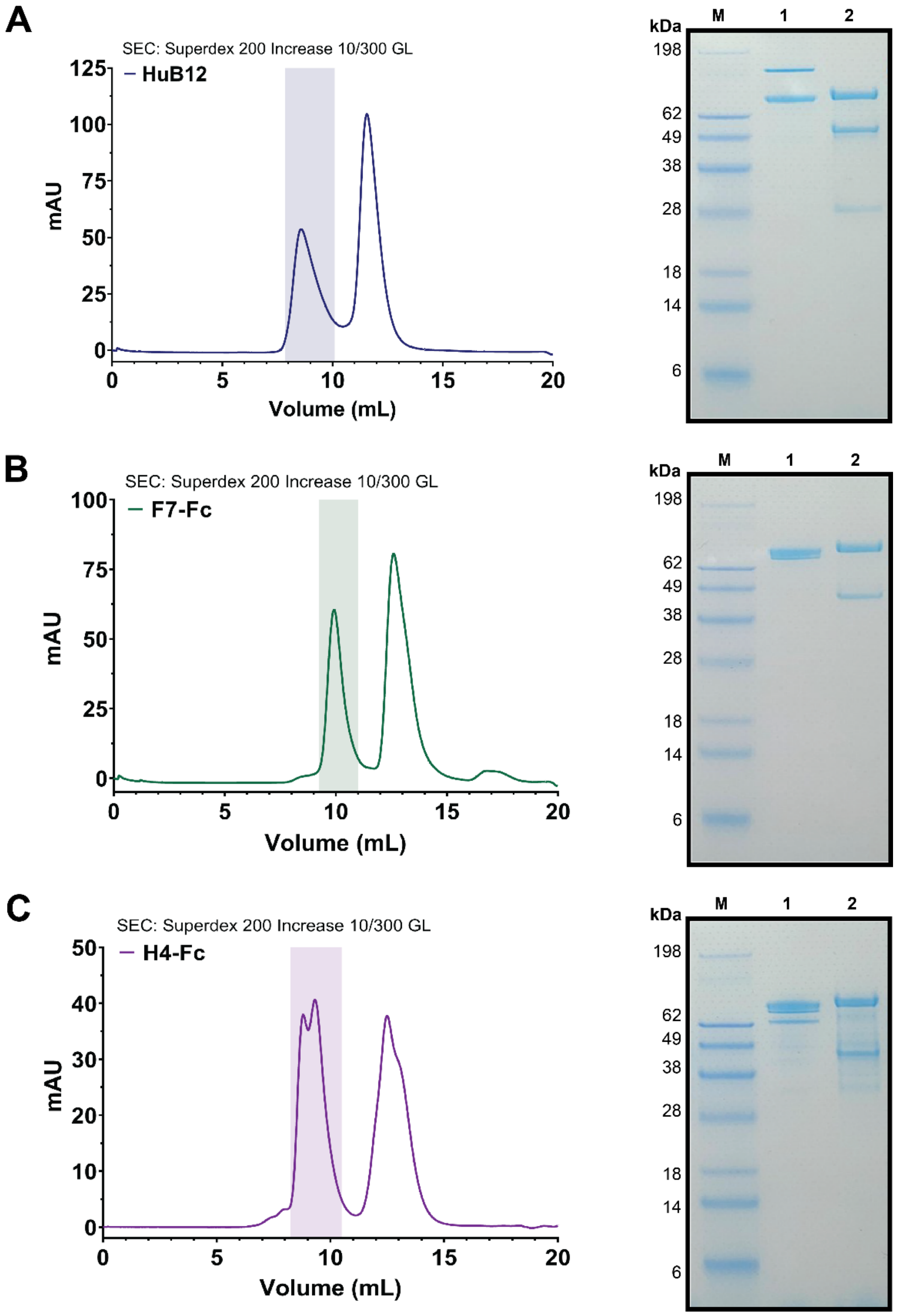


**Supplemental Fig. 9. Size-exclusion chromatography and SDS-PAGE FAP complexes.** Size-exclusion chromatography (left) of FAP in complex with huB12 (**A**), F7-Fc (**B**), or H4-Fc (**A**) with corresponding SDS-PAGE analyses of peak fractions (right) under non-reducing (1) and reducing conditions (2).

**Supplemental Fig. 10.**

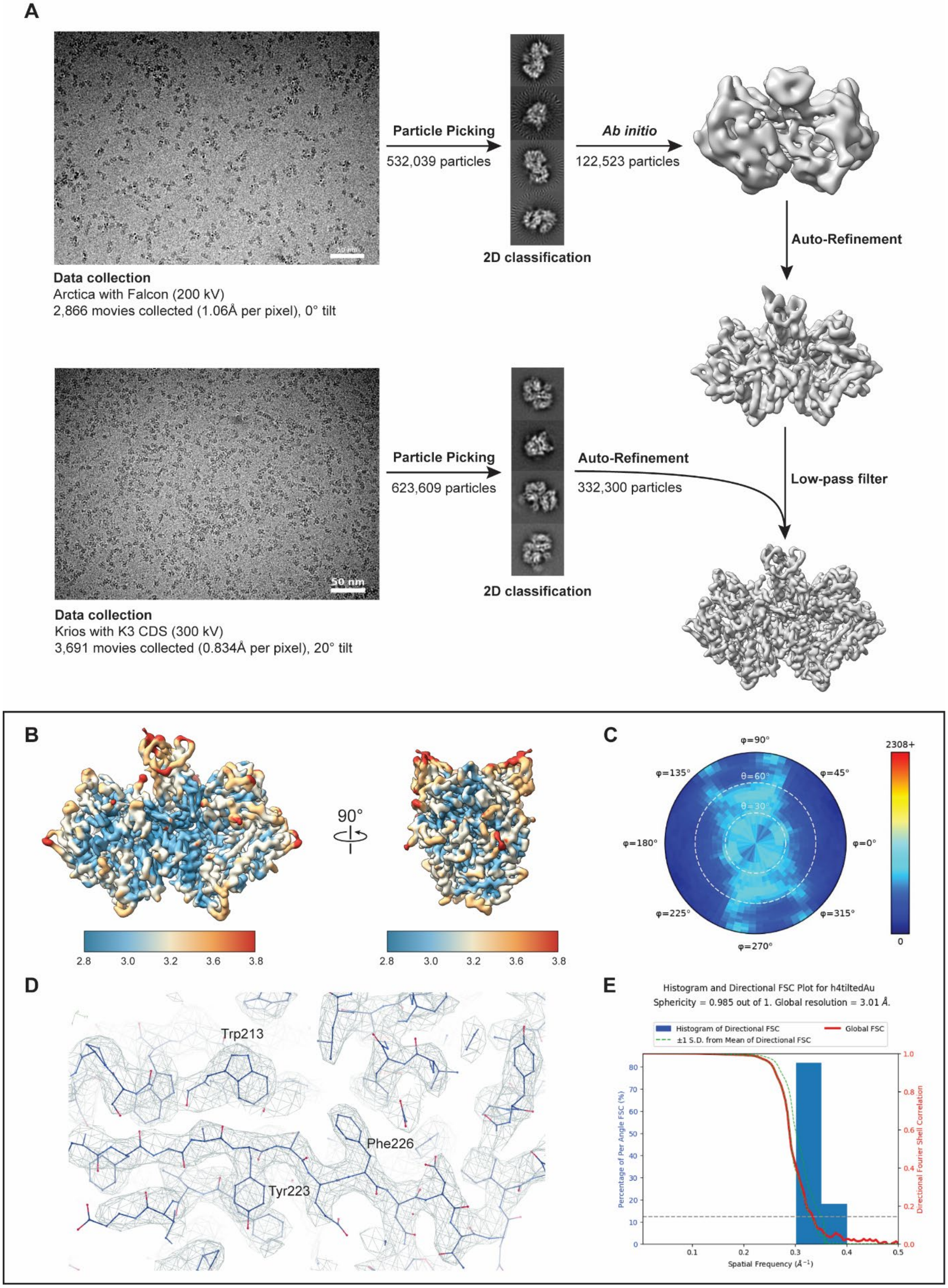

**Supplemental Fig. 10. Cryo-EM processing workflow and validation of FAP-H4.** (**A**) Representative micrographs, 2D class averages, and processing workflow for the FAP-H4 complex. (**B**) Local resolution estimates shown in two orthogonal views of the final reconstruction. (**C**) Angular distribution plot showing particle-view sampling in the final reconstruction. (**D**) Representative cryo-EM density with fitted atomic model, highlighting local map quality around selected aromatic residues. (**E**) All processing performed in *cisTEM*;[6] 3D Fourier shell correlation curve and directional FSC histogram for the final reconstruction.[7]

**Supplemental Fig. 11.**

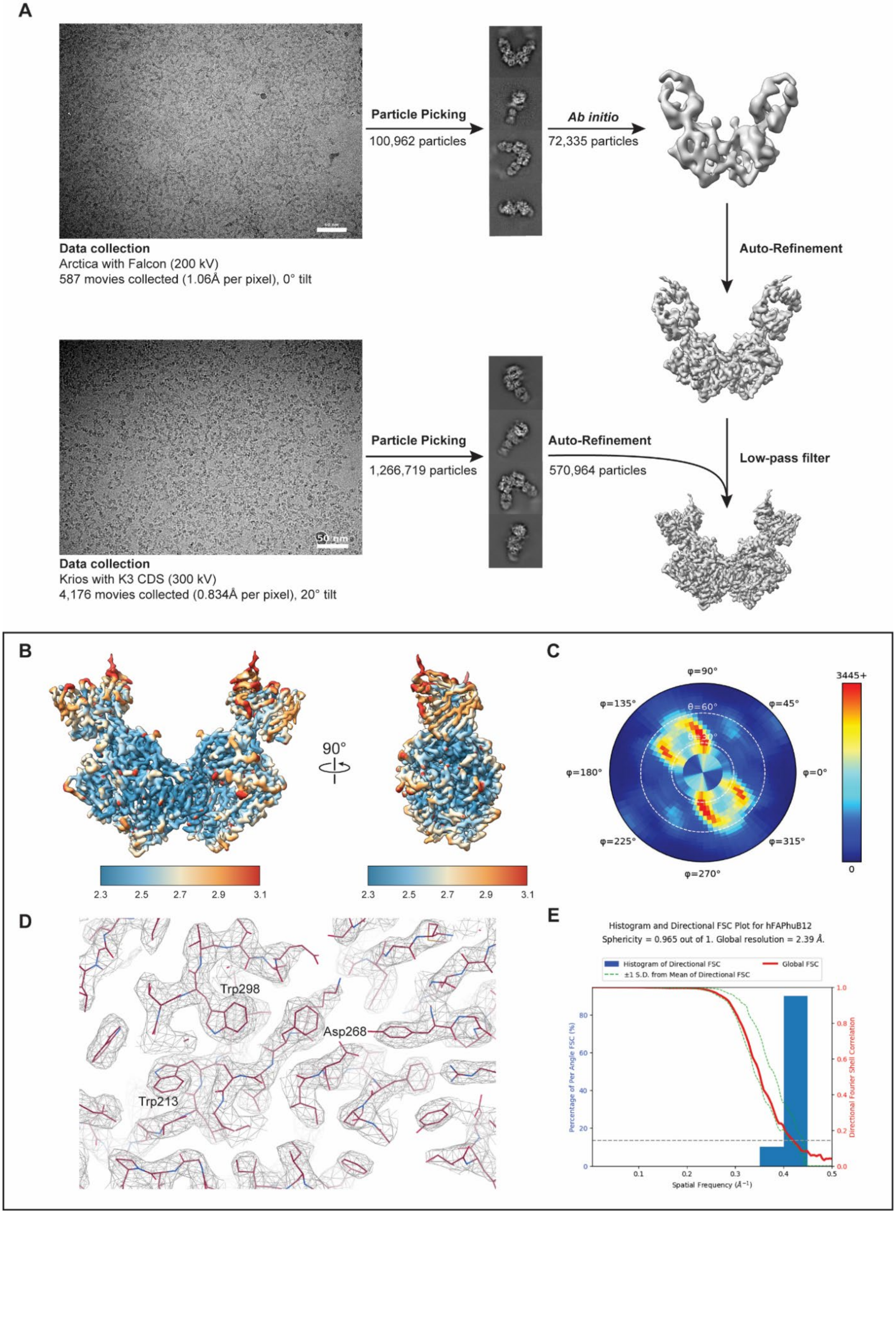

**Supplemental Fig. 11. Cryo-EM processing workflow and validation of FAP-huB12.** (**A**) Representative micrographs, 2D class averages, and processing workflow for the FAP-huB12 complex. (**B**) Local resolution estimates shown in two orthogonal views of the final reconstruction. (**C**) Angular distribution plot showing particle-view sampling in the final reconstruction. (**D**) Representative cryo-EM density with fitted atomic model, highlighting local map quality around selected aromatic residues. (**E**) All processing performed in *cisTEM*;[6] 3D Fourier shell correlation curve and directional FSC histogram for the final reconstruction.[7]

**Supplemental Fig. 12.**

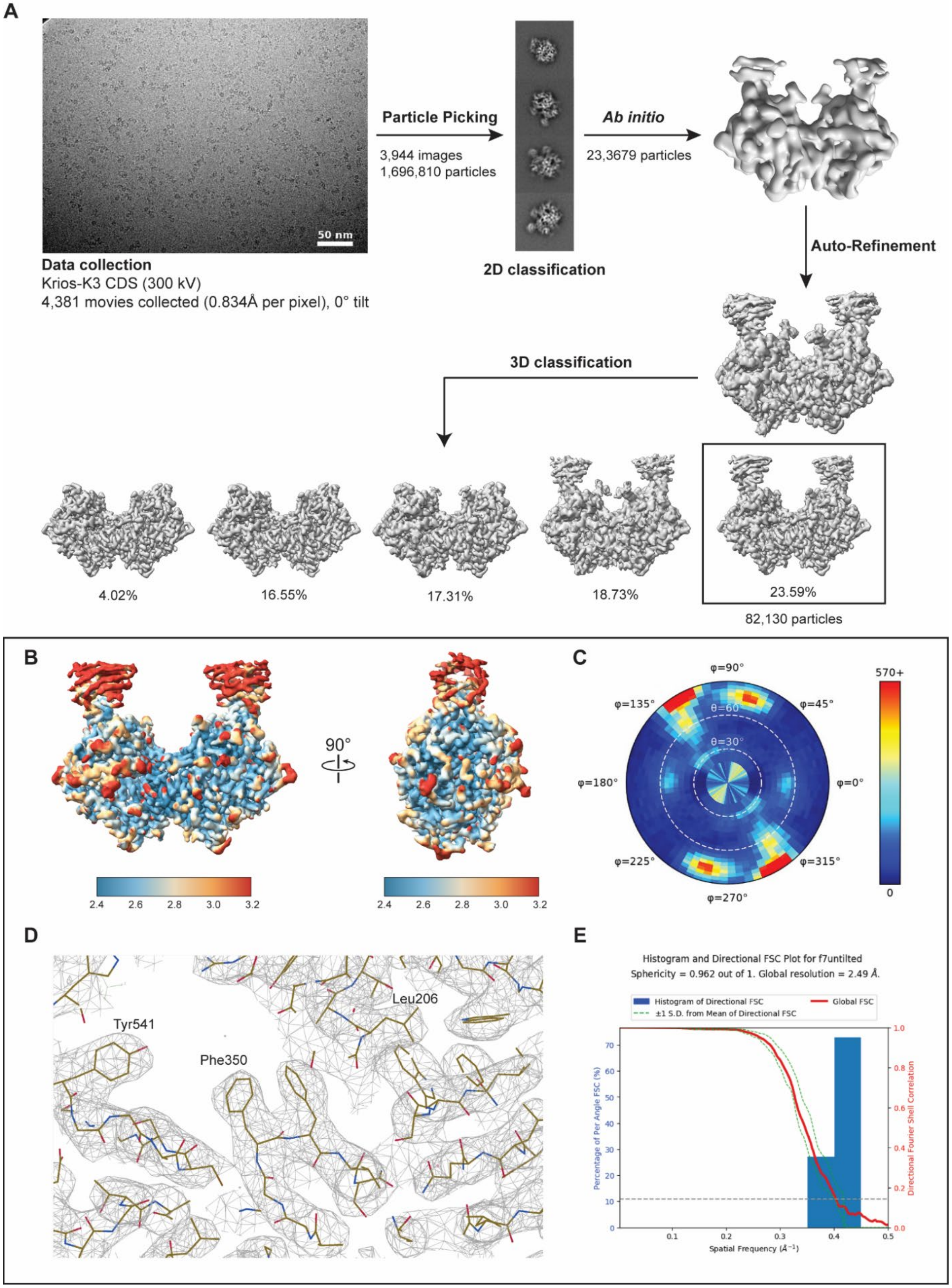

**Supplemental Fig. 12. Cryo-EM processing workflow and validation of the FAP-F7-Fc reconstruction.**
(**A**), Representative micrograph, 2D class averages, and processing workflow for the FAP-F7 complex. Particles were selected from 2D classification, used for ab initio model generation and auto-refinement, and further separated by 3D classification. The boxed 3D class was selected for final refinement. (**B**) Local resolution estimates for the final FAP-F7 reconstruction shown in two orthogonal views. (**C**), Angular distribution plot showing particle-view sampling. (**D**), Representative cryo-EM density with fitted atomic model, highlighting local map quality around selected residues. (**E**) All processing performed in *cisTEM*;[6] 3D Fourier shell correlation curve and directional FSC histogram for the final reconstruction.[7]

**Table 1. Cryo-EM data acquisition and model statistics.**

| Statistic | FAP-H4 (EMD-78501/PDB: 37US) | FAP-huB12 (EMD-78502, PDB: 37UT) | FAP-F7 (EMD-78503, PDB: 37UU) |
|---|---|---|---|
| **Data collection and processing** | | | |
| Microscope | Krios-G3 | Krios-G3 | Krios-G3 |
| Voltage (kV) | 300 | 300 | 300 |
| Detector | K3 (Counting) | K3 (Counting) | K3 (Counting) |
| Magnification (nominal/calibrated) | 105,000 | 105,000 | 105,000 |
| Data acquisition software | SerialEM | SerialEM | SerialEM |
| Electron exposure ($e^-/Å^2$) | 80 | 80 | 80 |
| Exposure rate ($e^-$/pixel/s) | 7.7 | 7.7 | 7.7 |
| Frames per micrograph | 80 | 80 | 80 |
| Pixel size (Å) | 0.834 | 0.834 | 0.834 |
| Defocus range (µm) | -0.5 to -2.0 | -0.5 to -2.0 | -0.5 to -2.0 |
| Stage tilt (°) | 20 | 20 | 0 |
| Micrographs collected (no.) | 3,691 | 4,176 | 4,381 |
| **Reconstruction** | | | |
| Image-processing package | *cis*TEM | *cis*TEM | *cis*TEM |
| Total extracted particles (no.) | 623,609 | 1,266,719 | 1,696,810 |
| Final particles (no.) | 332,300 | 570,964 | 82,130 |
| Symmetry imposed | C2 | C2 | C2 |
| Resolution (Å), FSC 0.143 (masked/unmasked) | 2.9/3.0 | 2.4/2.6 | 2.5/2.8 |
| 3DFSC sphericity | 0.985 | 0.965 | 0.962 |
| **Model composition** | | | |
| Protein residues | 1656 | 1882 | 1674 |
| Nucleotides | 0 | 0 | 0 |
| Ligands / glycans | 18 (NAG) | 20 (NAG) | 22 (NAG) |
| **Refinement** | | | |
| Refinement package | Phenix | Phenix | Phenix |
| CC (volume / mask) | 0.79/0.83 | 0.85/0.85 | 0.8/0.81 |
| Highest resolution used in refinement (Å) | 3.5 | 3.5 | 3.5 |
| **R.m.s. deviations** | | | |
| Bond lengths (Å) | 0.003 | 0.005 | 0.005 |
| Bond angles (°) | 0.874 | 0.971 | 0.977 |
| **Validation** | | | |
| Map-to-model FSC 0.5 (Å) | 3.1 | 3.0 | 3.5 |
| **Ramachandran plot** | | | |
| Outliers (%) | 0 | 0 | 0 |
| Allowed (%) | 1.76 | 1.71 | 1.80 |
| Favored (%) | 98.24 | 98.29 | 98.20 |
| MolProbity score | 1.37 | 1.32 | 1.43 |
| Poor rotamers (%) | 0 | 0 | 0 |
| Clashscore (all atoms) | 6.67 | 5.85 | 7.92 |
| C-beta deviations (%) | 0 | 0 | 0 |
| CaBLAM outliers (%) | 1.22 | 2.05 | 2.05 |

**Table 2. Sequence homology of human FAP (hFAP) across representative vertebrate species.**

| Species | hFAP Sequence Homology (%) |
|---|---|
| XP_013816149.1 *Apteryx mantelli* | 74 |
| XP_025950250.1 *Dromaius novaehollandiae* | 74 |
| XP_009674444.1 *Struthio camelus* | 74 |
| XP_009805927.1 *Gavia stellata* | 74 |
| XP_061860738.1 *Colius striatus* | 74 |
| XP_053926393.1 *Cuculus canorus* | 73 |
| XP_030310144.1 *Calypte anna* | 74 |
| XP_006037511.1 *Alligator sinensis* | 73 |
| XP_007421586.1 *Python bivittatus* | 72 |
| XP_020654035.2 *Pogona vitticeps* | 72 |
| XP_044274272.1 *Varanus komodoensis* | 70 |
| XP_031816435.1 *Sarcophilus harrisii* | 87 |
| XP_072468260.1 *Notamacropus eugenii* | 85 |
| NP_032012.1 *Mus musculus* | 90 |
| NP_620205.1 *Rattus norvegicus* | 89 |
| XP_040601389.1 *Mesocricetus auratus* | 90 |
| XP_016805441.1 *Pan troglodytes* | 100 |
| XP_004032755.2 *Gorilla gorilla gorilla* | 100 |
| NP_004451.2 *Homo Sapiens* | 100 |
| XP_003266222.1 *Nomascus leucogenys* | 99 |
| XP_003921979.1 *Saimiri boliviensis* | 97 |
| XP_073927060.1 *Castor canadensis* | 94 |
| NP_001091470.1 *Bos taurus* | 94 |
| XP_069430438.1 *Ovis canadensis* | 94 |
| XP_040085172.1 *Oryx dammah* | 94 |
| XP_019778827.2 *Tursiops truncatus* | 94 |
| XP_030708465.1 *Globicephala melas* | 94 |
| XP_007183240.3 *Balaenoptera acutorostrata* | 93 |
| XP_006196283.1 *Vicugna pacos* | 94 |
| XP_002924909.2 *Ailuropoda melanoleuca* | 94 |
| XP_027446978.1 *Zalophus californianus* | 94 |
| XP_005640309.2 *Canis lupus familiaris* | 93 |
| XP_011283649.1 *Felis catus* | 95 |
| XP_014587994.1 *Equus caballus* | 94 |
| XP_046515435.1 *Equus quagga* | 94 |
| XP_004375484.1 *Trichechus manatus latirostris* | 95 |
| XP_008256892.2 *Oryctolagus cuniculus* | 95 |
| XP_012615844.1 *Microcebus murinus* | 95 |

| | |
|---|---|
| XP_021121240.1 *Heterocephalus glaber* | 93 |
| XP_001512952.2 *Ornithorhynchus anatinus* | 84 |
| XP_069472422.1 *Ambystoma mexicanum* | 65 |
